# B-BIND: BIOPHYSICAL BAYESIAN INFERENCE FOR NEURODEGENERATIVE DYNAMICS

**DOI:** 10.1101/2024.06.10.597236

**Authors:** Anamika Agrawal, Victoria M. Rachleff, Kyle J. Travaglini, Shubhabrata Mukherjee, Paul K. Crane, Michael Hawrylycz, C. Dirk Keene, Ed Lein, Gonzalo E. Mena, Mariano I. Gabitto

**Affiliations:** Center for Data-Driven Discovery for Biology, Allen Institute; Human Cell Types Department, Allen Institute; Department of Statistics & Data Science, Carnegie Mellon University; Department of Laboratory Medicine and Pathology, University of Washington; Department of Medicine, University of Washington; Department of Physiology and Biophysics, University of Washington; Department of Statistics, University of Washington

**Keywords:** Hierarchical Bayesian Inference, Noisy Matrix factorization, Identifiable, Alzheimer’s Disease, Trajectory Inference

## Abstract

Throughout an organism’s life, a multitude of complex and interdependent biological systems transition through biophysical processes that serve as indicators of the underlying biological states. Inferring these latent, unobserved states is a goal of modern biology and neuroscience. However, in many experimental setups, we can at best obtain discrete snapshots of the system at different times and for different individuals. This challenge is particularly relevant in the study of Alzheimer’s Disease (AD) progression, where we observe the aggregation of pathology in brain donors, but the underlying disease state is unknown. This paper proposes a biophysically motivated Bayesian framework (B-BIND: Biophysical Bayesian Inference for Neurodegenerative Dynamics), where the disease state is modeled and continuously inferred from observed quantifications of multiple AD pathological proteins. Inspired by biophysical models, we describe pathological burden as an exponential process. The progression of AD is modeled by assigning a latent score, termed pseudotime, to each pathological state, creating a pseudotemporal order of donors based on their pathological burden. We study the theoretical properties of the model using linearization to reveal convergence and identifiability properties. We provide Markov chain Monte Carlo estimation algorithms, illustrating the effectiveness of our approach with multiple simulation studies across various data conditions. Applying this methodology to data from the Seattle Alzheimer’s Disease Brain Cell Atlas, we infer the pseudotime ordering of donors. Finally, we analyze the information within each pathological feature to refine the model, focusing on the most informative pathologies. This framework lays the groundwork for continuous pseudotime modeling in the analysis of neurodegenerative diseases.

## 1. Introduction

Modeling how cellular processes change in time due to endogenous or exogenous effects, in the context of discrete observation, is a fundamental problem in the biological sciences. Central to this problem is the longstanding goal of inferring a longitudinal trajectory delineating the dynamic evolution of each system. A wealth of research has cast this problem as learning a latent unobserved variable -termed *pseudotime*-dictating the progression of events where relevant measured quantitites are expressed as functions of this latent variable (Campbell and Yau, 2018, 2016; Hou et al., 2023; Street et al., 2018; Reid and Wernisch, 2016). Pseudotime inference was first motivated by the famous metaphor of Conrad Waddington in which undifferentiated cells progress along a continuum developmental trajectory to reach their committed cell fates (Waddington, 1940, 1957). Inference of such models is challenging for two reasons. Firstly, the lack of a universal definition of time renders the problem unidentifiable unless constraints are imposed. Secondly, and more critically, the successful inference of the parameters governing the system’s evolution requires the inference of the pseudotime variable as well, adding an additional layer of complexity. These are particular challenges in modern biological applications, where measurements describing progression can only be collected once per individual, providing only a cross-sectional snapshot of the system’s state. Aligning cross-sections along the pseudotime axis poses an additional inferential challenge, potentially impacting our ability to recover underlying dynamics in comparison with scenarios having multiple time measurements per individual or when pseudotimes are known beforehand.

Accurate temporal modeling is essential in the study of neurodegenerative diseases, in which we observe the aggregation of pathology in brain donors, but where the underlying disease state and its progression is unknown. Here, we focus on the study of neurodegeneration in Alzheimer’s Disease (AD) and consider biophysically-inspired models for protein pathology accumulation and the pseudo-temporal dynamics of diseases progression. Pseudo-time dynamics of disease progression and relevant parameters are inferred by using data from the recently released Seattle AD Brain Cell Atlas consortium (SEA-AD, https://portal.brain-map.org/explore/seattle-alzheimers-disease). Crucially, these cross-sectional measurements encompass a diverse set of pathophysiological markers that provide information on different stages of disease progression.

### 1.1. Relation to prior work

Pseudotime or ordering methods are widely applied due to their practical significance across various domains. While both share the goal of inferring data’s inherent order, pseudotime goes a step further than simple ordination by attributing significant importance to the pseudotime latent variable, quantifying the system’s progression along an inferred dynamic trajectory. Here, we offer a concise overview of related methods. In the analysis of cellular development using single-cell RNA-seq data, pseudotime is classically derived by computing a low-dimensional representation of each cell, inferring principal curves, followed by computing suitable minimum spanning trees (Magwene, Lizardi and Kim, 2003; Trapnell et al., 2014; Street et al., 2018). In contrast to these non-parametric methods, another line of methods frames the problem from a model-based, parametric perspective. For example, in the context of ecological niches and species modeling (Hui et al., 2015; O’Hara and van der Veen, 2024; Hui et al., 2023; Popovic, Hui and Warton, 2022; van der Veen et al., 2023; Roberts, 2020; Hoegh and Roberts, 2020), measurements of relative abundances of species are sorted along sites of greater abundance. There, generalized latent variable models are used to represent ordination as latent variables, and no true ordination occurs, but instead, multivariate latent variable ordination is used to represent underlying gradients that influenced species composition. In Political Science, the related problem of sorting latent preferences of agents (such as judges) given observed votes appears under the name of ‘ideal point’ models (Gelman and Hill, 2006) and this latent individual ordering is inferred by stating random-effects variables. In the field of psychometrics, models of changes in ability level over time utilize Item Response Theory (Kim and Camilli, 2014; Crane et al., 2006) to approximate non-linear growth.

Model-based approaches facilitate the simultaneous inference of the pseudotimes as well as any other relevant parameters. The method by Campbell and Yau (2018) models pseudo-time inference in the context of single-cell omics as a matrix factorization problem where parameters are inferred through Bayesian methods. Then, it is possible to characterize both the timing of events as well as the progression of each feature (genes). While their setup is like ours, their work focuses on modeling, and it doesn’t study in-depth the interplay of the different variables (such as the number of samples and dimension of cross-sections), the effects of pooling or shrinkage, and it omits rigorous consistency and performance guarantees as we do here.

Finally, a growing line of research applies pseudotime-based analysis to the context of aging (Pierson et al., 2019) and disease progression analysis, notably, to the study of cancer progression (Gupta and Bar-Joseph, 2008; Huang et al., 2023) and neural degeneration, including AD (Mukherjee et al., 2020). However, most studies fail to address a fundamental degeneracy present in latent variable inference problems, wherein pseudotime solutions can be scaled or affine transformed while still producing a valid solution—a mathematical indeterminacy inherent in many proposed frameworks. Our work attempts to resolve such degeneracy by imposing relevant mathematical constraints and is unique in that it addresses both the theoretical and applied aspects. We expect that this approach will help better inform the deployment of pseudotime inference methods for disease progression analysis.

### 1.2. Organization

This paper is organized as follows. In Section 2, we review the study of neurodegeneration and protein aggregation caused by AD pathology, and describe the data on which we based our work. In Section 3, we introduce a biophysically-inspired model to describe AD neuropathology aggregation. Biophysical considerations justify the use of generalized linear models with an exponential link function tying the unobserved timing of events to observations. We supplement this likelihood specification with priors that constrain the parameter space and represent complex dependency structures across measurements. We then obtain a family of Hierarchical Bayesian models that can be inferred using modern computational tools. In Section 4 we provide a mathematical analysis of a simplified linear model under which we can make precise statements regarding the identifiability, consistency, and convergence rates of our model as the number of individuals and measurements increase. Central to this analysis is the observation that the underlying inferential target can be understood at the most fundamental level as an instance of noisy matrix factorization. These theoretical results are supplemented with extensive simulations in Section 5, that show that in most aspects the theoretical results presented in the linear case also manifest in the generalized-linear setup. Simulations also show a few distinctive phenomena of generalized linear models that are not well captured by theoretical results on the linear case. In section 6, we apply our method to the SEA-AD dataset, and building on previous sections and specialized biological knowledge, we demonstrate that pseudotime and progression parameters can be reliably estimated. In Section 6.2, we extend our modeling framework to inform future experimental design. Finally, in Section 7, we discuss the significance and limitations of our work, and sketch future directions.

## 2. SEA-AD dataset

The present work is motivated by the need to understand the impact of pathological protein aggregation on cellular vulnerability across individuals spanning the complete spectrum of AD pathology. Quantitative neuropathological analysis was performed on the middle temporal gyrus (MTG) (Fig. 1A), a region of the cerebral cortex organized in a layered structure (from layers 1 to 6), of *d* = 84 postmortem brain donors included in the Seattle Alzheimer’s Disease Brain Cell Atlas (SEA-AD) study (Gabitto et al., 2023). Donors from the SEA-AD cohort were drawn from two ongoing longitudinal studies: the Adult Changes in Thought (ACT) and the University of Washington Alzheimer’s Disease Research Center (ADRC). We note that the SEA-AD cohort consists of donors with age distribution skewed towards advanced age (average age at death 88, SD=8) (Fig. 1B), which suggests a priori that high AD pathology values might be overrepresented in the cohort. The publicly available quantitative neuropathology dataset (accessible at sea-ad.org) comprises *M* = 35 total phenotypic measurements derived from 6 immunohistochemical stains targeting pathological proteins (hyperphosphorylated tau (pTau) and amyloid beta (A-*β*), along with additional markers selected for examining cellular changes in neurons, astrocytes, and microglia. Table 1 presents key markers used in this study and their corresponding extracted features. Immunostains for each protein and cellular marker are processed by a machine learning algorithm (https://indicalab.com/halo-ai/) that creates masks for each protein pathology, segments positive areas, and quantifies the burden of each pathology (Fig. 1A). Next, we provide a brief overview of general AD pathophysiology and elucidate how these pertinent variables contribute to understanding AD progression.

**Table 1.** Measurements used named by markers and the measured quantity

| Feature name | Biological measurement | Type of marker |
| --- | --- | --- |
| Percent 6e10 positive area | Amyloid-beta immunoreactive area | Pathological peptide |
| Number of 6e10 positive objects per area | Number of amyloid-beta plaques | Pathological peptide |
| Average 6e10 positive object area | Size of amyloid-beta plaques | Pathological peptide |
| Number of AT8 positive cells per area | Number of pTau positive cells | Pathological peptide |
| Percent AT8 positive area | pTau immunoreactive area | Pathological peptide |
| Average AT8 positive cell area | pTau immunoreactive area | Pathological peptide |
| Number of NeuN strong positive cells per area | NeuN immunoreactive area (mature neurons) | Cellular |
| Percent NeuN positive area | NeuN immunoreactive area (mature neurons) | Cellular |
| Percent of Iba1 and 6e10 positive colocalized objects | Iba1 immunoreactive area (microglia) | Cellular-pathological colocalization |
| Number of Iba1 and 6e10 positive colocalized objects per area | Iba1 immunoreactive area (microglia) | Cellular-pathological colocalization |
| Average Iba1 positive process area per cell | Iba1 immunoreactive area (microglia) | Cellular-pathological colocalization |
| Percent GFAP positive area | GFAP immunoreactive area (astrocytes) | Cellular marker |

**Fig 1.**
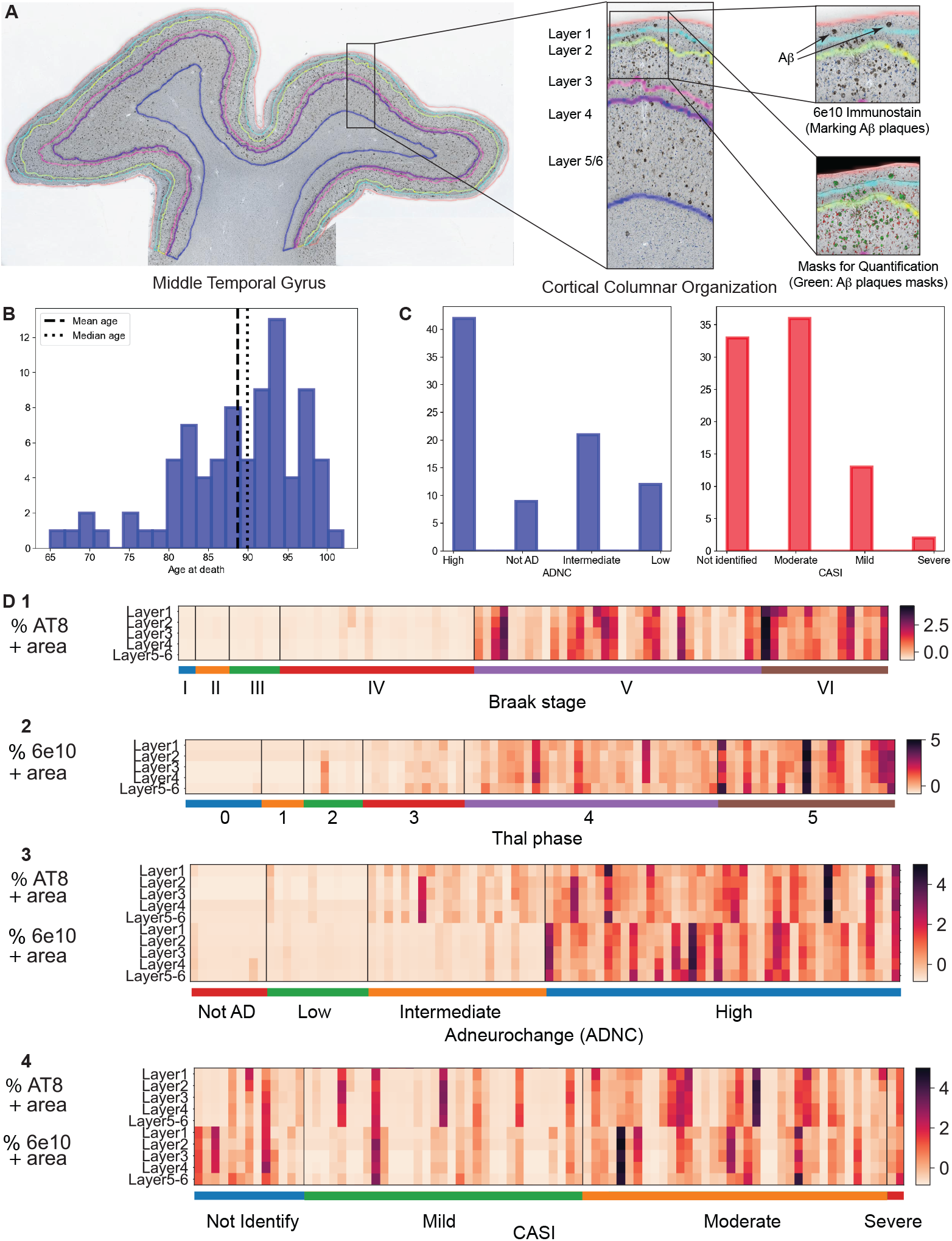
Seattle Alzheimer’s Disease Brain Cell Atlas (SEA-AD) data sets profiling neuropathology in AD brain donors. (A) Left, image of the middle temporal gyrus of a representative brain donor in which pathology data was stained and imaged over L = 5 cortical layers of MTG (middle box). Right, amyloid-beta aggregates stained with 6e10 stain are segmented by using ML algorithms (green masks) and quantified. (B) Histogram depicting age distribution of donors at death, average age at death = 88, median age = 90. (C) Histogram depicting the distribution of ADNC and CASI scores for SEA-AD donors ordered on the x-axis. (D1-D4) Neuropathological measurements, percentage AT8 (measuring phosphorylated Tau pathology) and 6e10 (measuring amyloid β pathology) positive stain area, for each MTG cortical layer organizing donors by (1) BRAAK stages, (2) THAL phases, (3) ADNC scores, and (4) CASI scores.

AD is the most common cause of age-related dementia and is characterized neuropathologically by the stereotyped spatiotemporal progression of two pathological proteins, intra-cellular hyperphosphorylated tau (pTau) in the form of neurofibrillary tangles (NFTs) and amyloid beta (A*β*) in the form of extracellular plaques. In AD, pTau and A*β* follow opposite progression patterns throughout the brain. Neuropathologists have mapped the progression of these pathological proteins and established discrete staging scales for each of them. These staging protocols order donors based on the presence or absence of pathological proteins in defined brain regions; Braak stages (I-VI) semi-quantitatively assess pTau NFTs (Braak and Braak, 1991), while Thal Phasing (1-5) semiquantitatively assesses A*β* plaques (Thal et al., 2002). A higher score on any scale indicates the presence of the relevant pathology across a greater number of brain regions. The AD neuropathologic change score (ADNC) (Hyman et al., 2012), attempts to integrate both scores into a general scale of AD progression. The two hallmark AD pathological proteins spatially overlap for the first time in the temporal lobe of the cerebral cortex, specifically the inferior and middle temporal gyrus (MTG). The MTG is thought to represent a key transition zone which delineates normal aging and preclinical AD from advanced stages of AD associated with widespread neocortical spread of pathology (Braak and Braak, 1991).

In addition to pathology measurements, the SEA-AD dataset encompasses a range of supplementary metadata pertaining to each brain donor, including clinical, cognitive, and demographic information. Established pathological assessments, previously described, such as Braak staging, Thal phasing, and ADNC are part of the repertoire of variables included, together with the last Cognitive Abilities Screening Instrument (CASI) (Teng et al., 1994a) test, administered during the donor’s lifetime to evaluate cognitive function (Fig. 1C). We used these per-donor-metrics to provide initial descriptions of the data and performed data-entry quality controls. Illustrative quantitative neuropathology measurements, categorized by staging and cognitive criteria, are depicted in Fig. 1D1-D4, which depicts a notable increment in pathological burden as metrics increase in value. While the full dataset consists of *M* = 35 pathological markers, we use a subset of those in our model. We considered *M* = 12 markers (described in Table 1), removing noisy biomarkers that have low signal-to-noise ratio or that were sparse, i.e. data was available just for a few donors.

## 3. Model Description

We aim to elucidate the challenge of modeling the inherent variability in a biological process by using a latent variable that represents pseudotime. The latent variable is named pseudotime because it not only orders observations but also governs the progression of the biological process. While we specifically focus on AD pathology aggregation (see Section 2), this framework has broader applications, such as investigating aging-related phenotypes or studying different cell types in cellular development.

In modeling AD progression, we assume donors traverse a single disease trajectory spanning the entire disease spectrum, from no disease to the end stage, yet this trajectory remains unknown. Donors are randomly selected across the disease spectrum, driven in practice by donor availability, and we lack prior knowledge of the sampling distribution, *p*(*t*). For each of the donors, we collect multiple measurements reflecting the state along the disease trajectory, including varying levels of pathological protein aggregates. Lastly, observations are corrupted by observational noise (Fig. 2). Notable previous work in aging studies has observed that to ensure identifiability of the link function between the latent pseudotime and the observational process (represented by *f* (*t*) in Fig. 2) should be monotonic and injective (Pierson et al., 2019). However, in their work, time was a known variable, compared to our case in which nothing is known about latent pseudotime.

**Fig 2.**
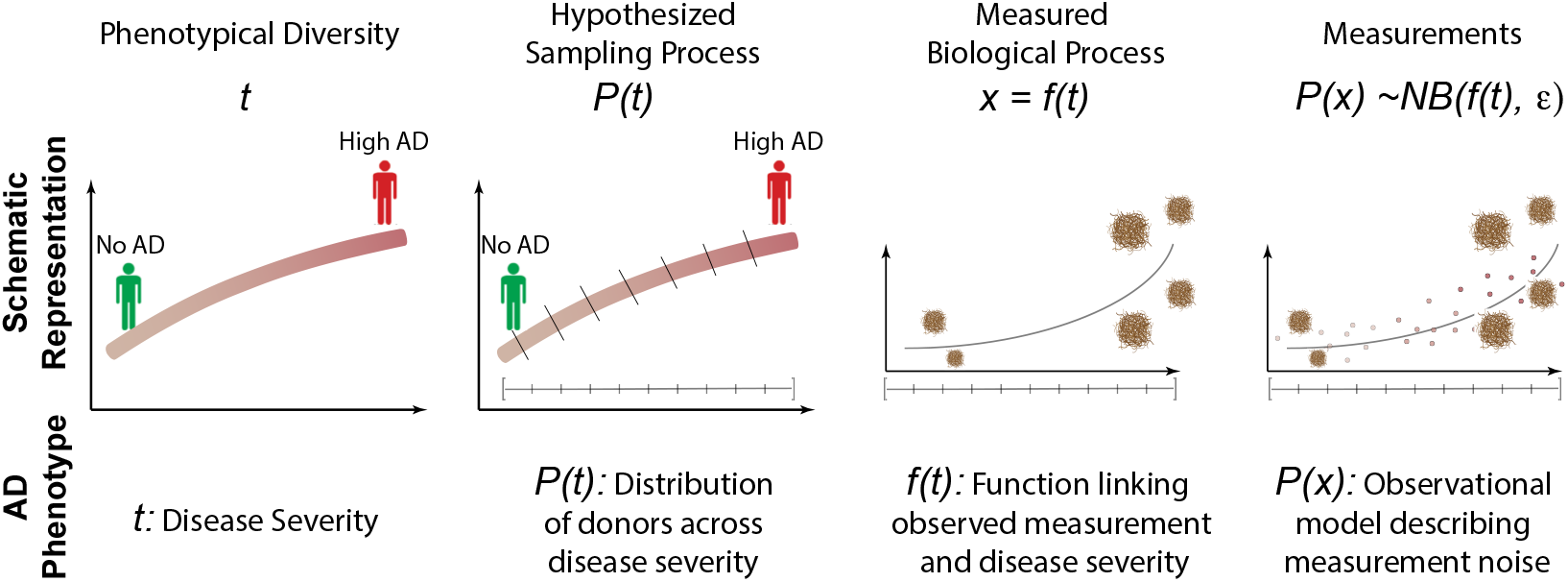
Schematic representing the modeling of a general biological process by a latent variable, t, termed pseudotime. In this work, pseudotime represents a scale that describes the trajectory of Alzheimer’s Disease along the entire disease spectrum. Donors are sampled along AD trajectory from distribution p(t). Then, multiple observed biophysical processes x are measured from each donor. These measurements can be linked to the underlying pseudotime by a mean function, f (t), and when observed, are corrupted by noise (described by the distribution p(x|t) of observational noise.

In the next sections we construct a model for cross-sectional SEA-AD donor data, 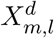, comprising quantitative neuropathological measurements from *d* = 1 … *D* = 84 donors, en-compassing *m* = 1 … *M* = 12 distinct protein measurements taken across *l* = 1 … *L* = 5 cortical layers in MTG. These measurements reflect the accumulation of various pathological proteins within each donor. We define a donor-specific one-dimensional latent variable, referred to as *t*_*d*_ ∈ [0, 1]. This variable represents the pseudo-temporal score that organizes donors according to their varying disease burdens, from low (*t*_*d*_ = 1) to high (*t*_*d*_ = 1). Inspired by biophysical principles of protein aggregation, and the cited aging-related studies (Pierson et al., 2019), we will find that a monotonic and injective exponential link function best accommodates our data. We will infer all model parameters through Bayesian inference.

### 3.1. Deterministic Biophysical model of Pathology Aggregation

Alzheimer’s Disease is characterized by the proliferation of misfolded protein aggregates. As mentioned before, amyloid-Beta (A*β*) aggregates and Tau neurofibrillary tangles (*τ* -NFT) are two examples of well-studied protein aggregates that have been widely used as pathological markers to study the spread and severity of AD (Braak and Braak, 1991; Thal et al., 2002). Several mechanistic models, chiefly rooted in polymer kinetics, have been used to study the aggregation and seeding dynamics of protein aggregates (Vaquer-Alicea and Diamond, 2019; Davis and Sindi, 2016). In our work, we will leverage the Nucleated Polymerization Model (NPM), which is the most widely accepted model used to describe protein aggregate propagation dynamics (Masel, Jansen and Nowak, 1999), to describe the temporal progression of AD-associated pathology markers.

The NPM model describes a cycle of aggregate growth followed by multiplication, leading to the dominance and growth of toxic protein aggregates. The growth of such aggregates occurs through attachment with surrounding ‘healthy’ monomeric units, which is then followed by a fragmentation process that multiplies the number of ‘seeds’ capable of growing into many more misfolded aggregates (Fig. 3A). Once the monomers are incorporated, they become a part of the “unhealthy” or misfolded population. The misfolded aggregates also tend to be more stable than their healthy counterparts, such as in the case of prionic proteins (Tompa et al., 2001). In an aggregation model for a spatially limited domain (e.g. a single anatomical region of the brain), the important dynamical parameters to consider are the initial seeding levels, denoted by *β*_0_, and the rate of multiplication through aggregation and division, denoted by *β*_1_ ((Fig. 3B). In the ‘early’ time regime, i.e. when the available pool of healthy protein monomers is not limiting, we will describe the evolution of a pathological protein *X*_*m,l*_ for measurement *m* in layer *l* with the following dynamics arising from the NPM biophysical model (Masel, Jansen and Nowak, 1999):

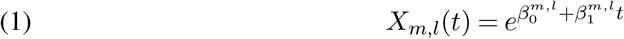

where *t* denotes the latent pseudotemporal variable describing the progression of AD severity. The details of the kinetic model of pathology propagation, and the simplifying assumptions used to arrive at such functional form, are included in Supplementary Material, Appendix Section A.

**Fig 3.**
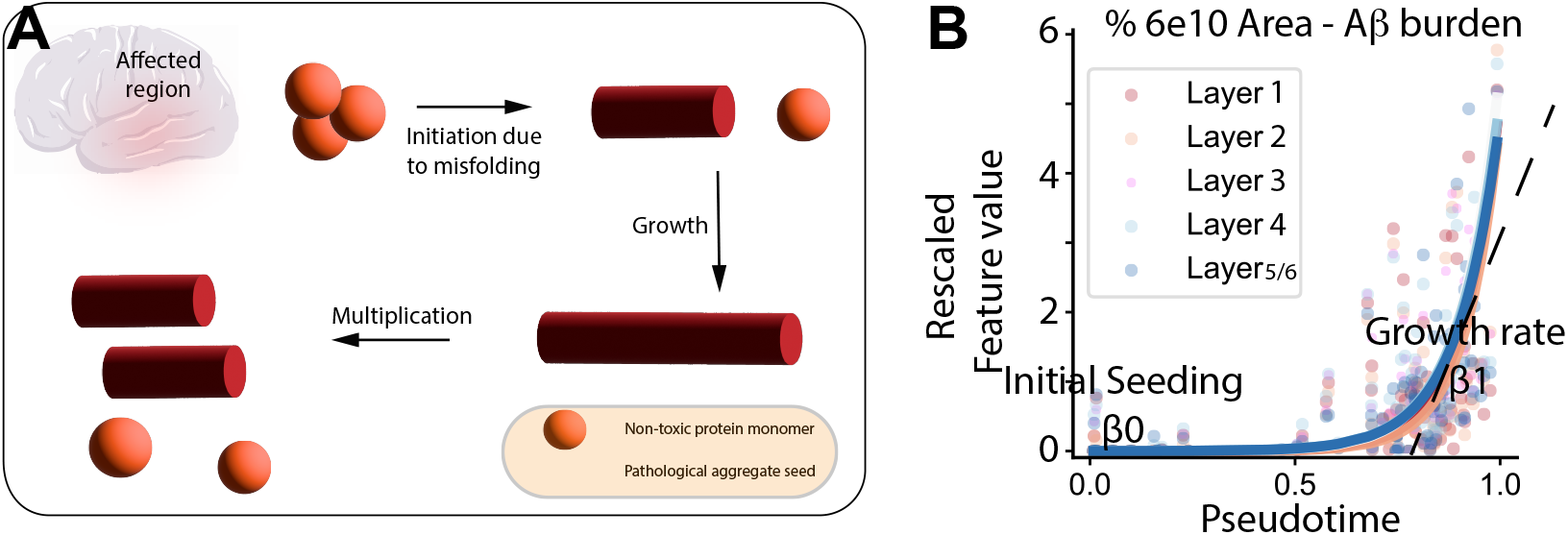
We assume that AD protein pathology aggregation can be described by an exponential process. A) Schematic representing protein aggregation process. Due to protein misfolding, an initial seeding event is triggered which later leads to aggregate growth. Randomly, protein aggregates break down multiplying in number. The conjunction of growth and multiplication events can be described by exponential dynamics. B) Scatterplot depicting measurement for Aβ burden, the percentage area stained positive by 6e10 antibody, in each cortical layer ordered according to our model. For each protein pathology, the model fits an initial seeding variable, β_0_, and a growth aggregation rate β_1_.

### 3.2. Modeling the Generative Distribution of Observed Pathology

The above biophysical argument suggests to consider a trajectory that is an exponential transformation of an affine function of the latent pseudotime variable. To account for observation noise, we couple this latent trajectory with a sampling model where observed counts 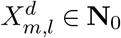 describ-ing pathological proteins follow a negative binomial distribution. Specifically, we consider a count parameter *A*_*m,l*_ ∈ **N**^+^, and a probability of success in the (negative) logit scale *ρ* ∈ **R** such that the likelihood of 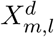 given *A*_*m,l*_ and 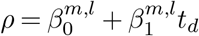 is given as

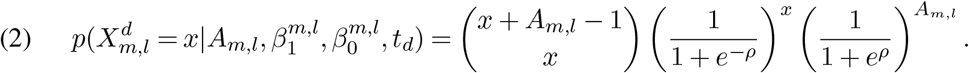

The mean and variance of the above distribution are given by 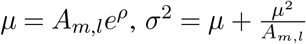.This implies that the conditional mean of 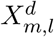 given 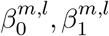 and *t*_*d*_ is exactly the quantity *X*_*m,l*_(*t*) defined in Eq. (1), up to the *A*_*m,l*_ factor that can be absorbed into 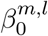. Consequently, the negative binomial model can be understood as providing a noisy version of the underlying dynamics. As customary, we understand *A* as an overdispersion parameter *A* (Gelman, Hill and Vehtari, 2021) modulating different levels of observation noise whereby in the limit *A* →∞ we recover the usual Poisson regression. Eq. (2) is one instance of a generalized linear model with a logarithmic link function. Still, other choices could be made if observations were of a different nature. For example, logistic regression could be used if observations were binary.

#### 3.2.1. Priors

In addition to the likelihood in Eq. (2), we must specify priors for the model parameters Θ = *{t, β*_0_, *β*_1_, *A}*. First, we specify a sequence of independent donor-specific pseudotimes following a Beta distribution with parameters *t*^*a*^, *t*^*b*^:

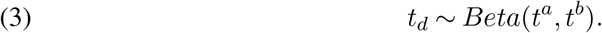

In particular, *t*_*d*_ ∈ [0, 1]. The decision to choose such a bounded interval is two-fold. At the conceptual level, biological systems are constrained in space and time. This implies in our application to AD that protein propagation and disease trajectory do not last indefinitely. When looking at a particular brain region, the accumulation of pathological proteins will range from 0 to a high number, with the last case implying that the pathological protein has covered the entire region. At the mathematical level, as we will later show in Section 4, choosing a bounded interval enables identifiability of the entire system, a desired property in our setup where our latent variable represents not only ordination but also pseudo time. The case *t*^*a*^ = *t*^*b*^ = 1 corresponds to a uniform prior. Although this prior may seem unbiased, it encodes strong beliefs about the phenotypical diversity of the sample; for example, that the average severity level is 0.5. This justifies the use of a more general *Beta* prior to encode information on the sampling process. We will see in Section 4 that we can benefit from suitable choices of this prior.

We must also specify priors on 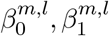 parameters that represent the baseline level of pathology, and the exponential rate of pathological growth of each of the markers. We will consider two models, one with measurement and layer-independent priors on 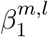 and the second with hierarchical priors pooling information across layers. In the former case, we are assuming that only the protein aggregation rate (through its parameter *β*_1_) can be related across layers, and for each measurement, we will infer a covariance matrix across layers. Finally, we consider priors for the over-dispersion parameters *A*^*m,l*^ since they are not revealed to us. In this case, we impose half-normal distributions.

In summary, the plate diagrams of our hierarchical and fully factorized models are depicted in Fig. 4, and we detail next the entire generative process of our model:

**Fig 4.**
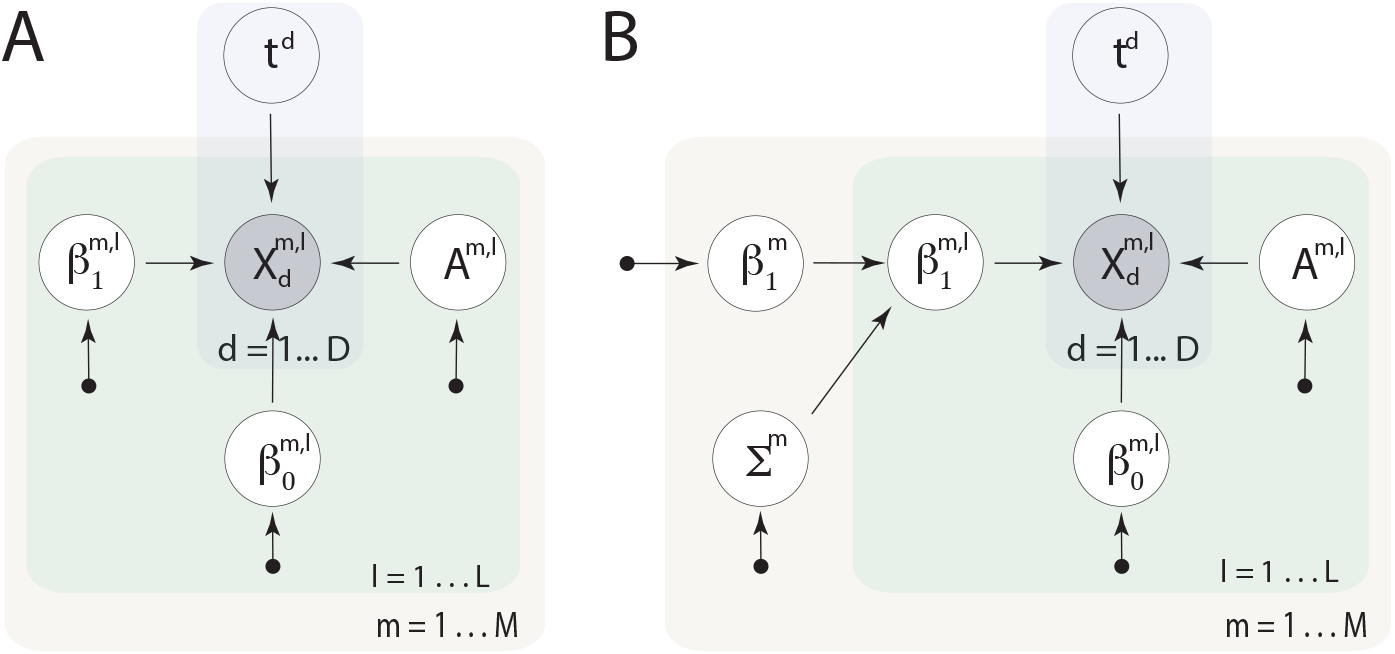
Plate diagram depicting a model in which parameters are inferred independently in each measurement and each layer. **(A), and model with hierarchical priors pooling information across layers in each measurement (B)**. In both models, pathological measurements are depicted by 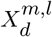 where m = 1 … M index each of the measurements and l = 1 … L each layer. The latent pseudotime variable is denoted by t_d_, where d = 1 … D index the brain donor. Measurement and layer-specific initial-seeding, growth-rate, and total-counts parameters are denoted by 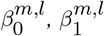 and A_m,l_. In the case of the hierarchical model in (B), we pool information across layers and infer covariance matrices Σ_m_, assuming that each layer has its own initial seeding events but their growth rates are similar.

1. For each donor, *d*
  a. Choose *t*_*d*_ ∼ Beta(1, 1)
2. For each measurement, *m*
  a. **If** Hierarchical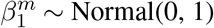 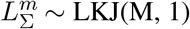 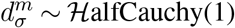 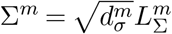 **Else** Pass
3. For each layer, *l*
  i. Choose 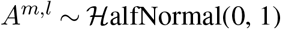
  ii. Choose 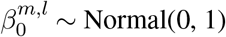
  iii. **If** Hierarchical **Then** Choose 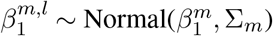 **Else** Choose 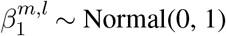

### 3.3. Posterior Bayesian Inference through HMC sampling

To perform posterior inference from our models we used numpyro version 0.13.0 (Bingham et al. (2019); Phan, Pradhan and Jankowiak (2019)), a Python based probabilistic programming environment. Similar to the more traditional probabilistic language, Stan (Carpenter et al. (2017)), numpyro is equipped with effective posterior sampling algorithms such as Hamiltonian Monte Carlo (HMC, Duane et al. (1987); Neal et al. (2011)). In particular, we utilized the implementation of the No-U-Turn Sampler (NUTS) (Hoffman et al. (2014)) kernel to obtain posterior samples, and monitored convergence using R-hat and effective sample size metrics (Gelman et al. (2021)).

To infer the posterior parameters of our models (Fig. 4) using the SEA-AD dataset, we sampled from the posterior with 5 HMC chains run sequentially for 5000 warm-up iterations, and then 1000 samples were collected in each chain. We randomly rotate the seed for the pseudo-random number generator in each chain. To ensure convergence, we discarded the chains that had an R-hat below 1.05 or had an effective sample size below 50. The results from these fits are described in Section 6. For experiments with synthetic data, we used different parameters to the ones described above (see Supplement Section on Proofs for details).

## 4. Model analysis

Our problem departs from the classical regression setup in that we not only estimate the regression parameters *β* but also must infer the latent parameter *t*. Here, we delve into the inferential implications of the joint estimation of *t* and *β*; we study the fundamental statistical limitations to our problem and argue that our modeling choices are well-justified in that they enable efficient use of all available sources of information to counter these limitations.

Results in this section are statements regarding conditions for identifiability and the best possible convergence rates for the mean squared error (MSE) of estimators of *β* and *t*. Although we keep our Bayesian estimators in mind, this analysis is rather concerned with the properties of the problem itself than with our particular method. We will later show in Section 5 that our Bayesian estimators (understood as posterior means) exhibit the behavior of the best possible estimators.

Instead of the generalized linear model in Eq. (2), here we consider the simpler linear model where a rich theoretical understanding of statistical limits is readily available:

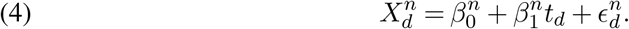

We understand *X* as a *D × N* matrix by vectorizing layer and feature information and treating them as a unique column ‘feature’ indexed by *n* = 1, …, *N* where *N* = *L × M*. Here, 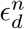 are i.i.d. Gaussian centered variables with variance *ϕ*^2^. These noise variances play an analog role to the overdispersion parameter *A* in Eq. (2).

The key observation is that Eq. (4) corresponds to a rank-two noisy factorization problem, so that at the most elementary level, the estimation limits of *β* and *t* are the limits of this low-rank factorization. Although the theory is general enough to enable the analysis of (4), for simplicity we will neglect the 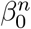 term and study the rank-one noise matrix factorization

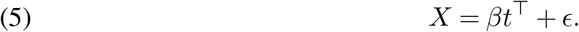

To apply existing statistical results, we divide the estimation of *t* and *β* into the estimation of the normalized parameters *β*^*′*^ = *β/*||*β*||_2_, *t*^*′*^ = *t/*||*t*||_2_ and the individual signal strengths 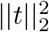 and ||*β*||^2^. To state consistency results, we conceive limiting experiments consisting of sequences of true *{t}*_*d*_, *{β}*_*n*_ (for example, obtained by sampling from their population distributions), and of matrices *X* and *ϵ* that are revealed as either *D* or *N* (or both) get larger. For our asymptotic statements on *D* or *N* we must assume that 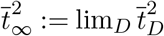 and 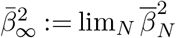 exist, where 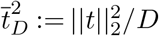 and 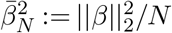,respectively.

### 4.1. Subspace estimation

We first study the ability to estimate the normalized parameters *t*^*′*^ and *β*^*′*^. We use the standard Frobenius metric for comparing subspaces *U*_1_ and *U*_2_(see Appendix for details), which in the one-dimensional case reduces to

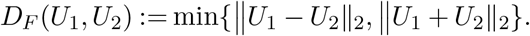

To obtain estimation bounds, we turn to specialized, best-known risk analysis for subspace estimation Cai and Zhang (2018) based on variants of the well-known Wedin’s sin Θ perturbation theorem Wedin (1972). Let 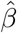 and 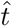 be the first left and right singular vectors of *X* (in particular, they have unit norm).

#### Proposition 1.

*As a consequence of Theorems 3 and 4 in Cai and Zhang (2018), we have*

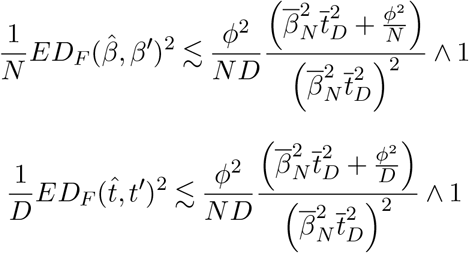

*These estimators are rate-optimal in the sense that the risk of any other estimators of β*^*′*^ *and t*^*′*^ *have risk lower bounded by the right sides above (up to constants)*.

We can use Proposition 1 to understand how estimation performance for *β*^*′*^ and *t*^*′*^ improve as we increase either *D* or *N* (or both). We observe that for fixed *N*, the average MSE in the estimation of *β*^*′*^ decreases at the rate *D* if 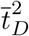 is assumed to remain bounded away from zero (we are excluding sparse non-informative sequences of donors), which is the same rate that would be achieved if *D* were known. Therefore, adding more donors is helpful in the estimation of *β*^*′*^. Likewise, if *D* is fixed, then the average MSE over *t*^*′*^ decreases at the rate 1*/N* if 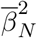 (i.e. we are not adding too many non-informative features), implying that adding more features help. Also, larger magnitudes 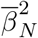 are beneficial for the estimation of *t*^*′*^ and larger magnitudes of 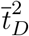 are beneficial for the estimation of *β*^*′*^.

### 4.2. Signal strength estimation

Proposition 1 suggests that the spaces spanned by *t*^*′*^ and *β*^*′*^ can be estimated as easily as in the usual regression case (*t* known). We now turn to the problem of estimating the signal strengths ||*β*|| and ||*t*||. The ability to estimate these parameters consistently is critical since otherwise, we are at risk of inflating the significance of our estimators of *β*: for example, if ||*β*|| is grossly overestimated, we may come to falsely detect many *β*_*n*_ as significant features explaining the severity of the pathology.

Estimating the signal strengths is more delicate since identifiability issues might be at play. The main result is that by imposing the constraint that severity levels belong to [0, 1] (or any other compact interval defined beforehand), we can identify ||*β*|| and ||*t*|| if both *N* and *D* simultaneously grow to infinity.

#### Proposition 2.

*Let t*_*d*_ *be sampled from a distribution* **t** *supported on* [0, 1]. *Suppose that both t*_∞_ *and β*_∞_ *exist. Then, there are estimators of* 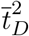 *and* 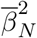 *converging almost surely to* 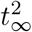 *and* 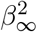 *as N and D grow to infinity*.

The mathematical details are presented in the Appendix. At a high level, the proof consists of first showing that we can first identify the product *β*_∞_ *× t*_∞_. Then, we can use the information on the support of **t** to re-scale the principal vector 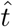 so that it will lie in [0, 1]. This re-scaling enables the identifiability of each of the terms *β*_∞_ and *t*_∞_. The statement in Proposition (2) holds whether or not *ϕ*^2^ is known: in the latter case, we can estimate it incurring an asymptotically negligible bias.

Proposition 2 gives the optimistic result that the entire system can be identified, but the main downside is that we are forced to consider simultaneous asymptotics in *N* and *D*. Unfortunately, as we will show, this heavy requirement cannot be relaxed, and the estimation of ||*β*|| will be biased. In other words, the estimation of ||*β*|| will be biased even if a cohort of infinite donors were available. We will comment on strategies to overcome this limitation. Even if these biases persist, the overall message of Proposition 2 is that if multiple informative features *β* are available, they should all be used to inform inferences.

## 5. Simulation studies

The previous results provide us with tools to reason about inferential capabilities in our model. In this section, we complement these results with a comprehensive set of simulations. These simulations not only give an empirical validation to the theory in Section 4; more importantly, we use them to understand our procedure’s behavior in cases not covered by the theory.

This gap arises in three aspects: First, results in Section 4 are concerned with the estimation problem and not with our particular estimator. Second, they concern a simplified model that doesn’t incorporate the effect of a nonlinear link function and a non-Gaussian likelihood, and where coefficients *β* are not assumed to possess a hierarchical structure that we could possibly benefit from. From the experiments in this section, we can make the following claims

- Our Bayesian estimator exhibits the behavior anticipated by Propositions 1 and 2 in that simplified setup (Sections 5.1, 5.2)
- The use of suitable priors on *t* can be helpful to overcome the identifiability issues described in Section 4 (Section 5.1)
- Beyond the simplified linear model, the qualitative behavior of estimators still matches the phenomena described in the previous section, although some additional new phenomena emerge:
  – In non-linear/non-Gaussian models, hierarchical models for *β* are beneficial if such structure is present in the data. We don’t observe a benefit from using hierarchical models in linear Gaussian models (Section 5.3)
  – Non-linearities in the link function imply that different features can be differentially informative at different levels of severity. This implies the need for careful choice of features and donor experimental design (Section 5.4)

Here, we only describe the main features of each experiment but defer a more comprehensive explanation of each setup and parameter and sampler values to the Appendix.

### 5.1. Elementary simulations in the linear model

We start by studying the empirical performance of our Bayesian estimators in the setup of the linear model in Eq. (4). Unlike the full hierarchical model of Section 3, we consider a flat prior over *β*. We study different choices of true pseudotime distribution **t** and pseudotime priors *p*(*t*). The most elementary choice on pseudotimes *t*_*d*_ consists of them being sampled equispaced in [0, 1]. We also consider sampling from a *Beta*(0.2, 0.2) distribution and from a “cubic” distribution; i.e., where **t** is proportional to *t*^3^ on [0, 1]. As priors we consider the uniform distribution in [0, 1] (representing the equispaced sampling) and the *Beta*(0.2, 0.2).

We study the performance in estimation of *β* comparing against the regression estimator that would be obtained if the pseudotimes were known. If that was the case, *β* = (*β*_0_, *β*_1_) could be inferred via least squares with closed-form expressions.

As shown in Fig. 5 and Supplementary Fig. S4, pseudotime inference signifies a sizable increase in the MSE of the posterior mean of *β* over the baseline where pseudotimes are known. As expected, the MSE decreases as the variance parameter *ϕ*^2^ increases (here, assumed known). Proposition 2 anticipates that *β* may be biased even if *D* is arbitrarily large. This is consistent with Fig. 5A, where the MSE of *β* doesn’t converge to the baseline, indicating this persisting bias. Although uncertainty quantification is not the main intent of our work, we show that similar behavior occurs when we compare the posterior variance of our estimator with the posterior variance of the usual linear regression estimator (Supplementary Fig. S4).

**Fig 5.**
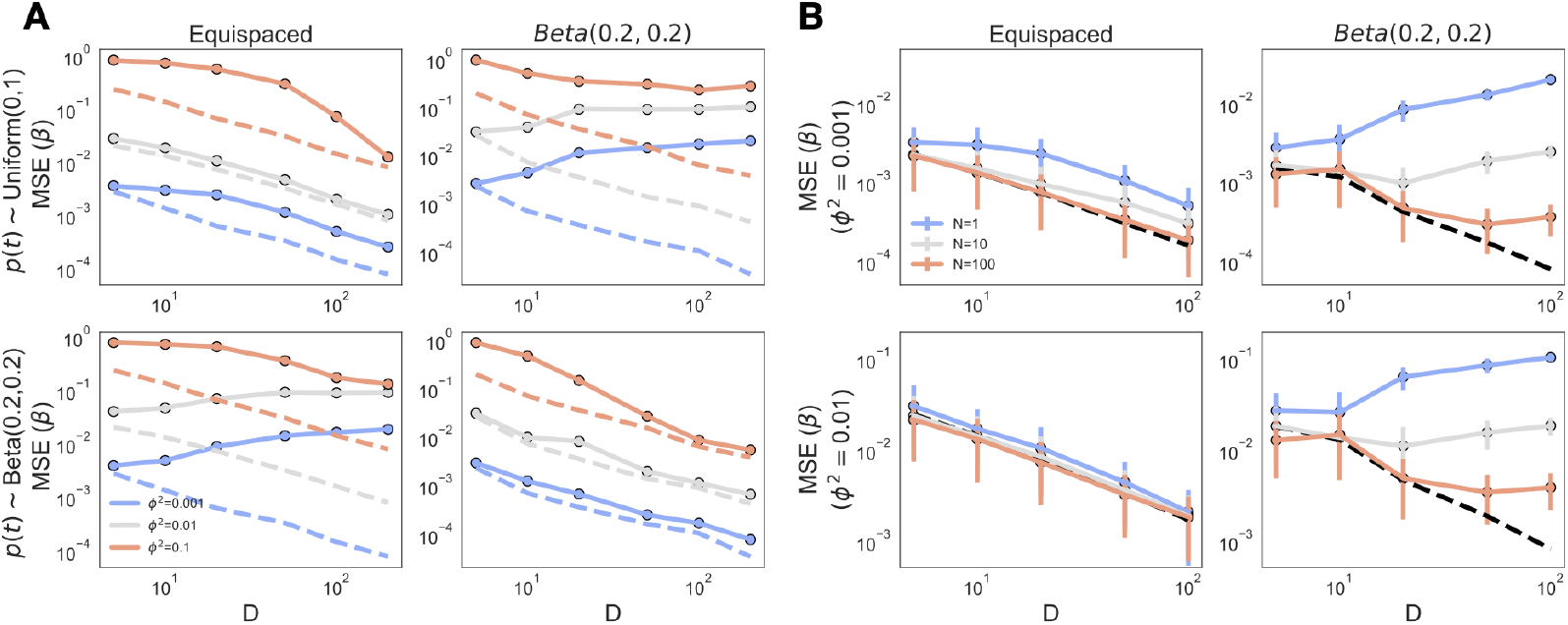
Effect of choices of pseudotime priors. **(A) number of features** N **(B) and in the linear model (Eq. on estimation error (MSE) of** β = (β_0_, β_1_). (A): Priors p(t) ∼ Uniform(0, 1) and p(t) ∼ Beta(0.1, 0.1) are shown on different rows and population distribution of pseudotimes **t** on different columns. Colors indicate noise levels ϕ_2_. We used a single feature for this experiment (N = 1). (B): Different rows correspond to different observation noise levels ϕ_2_, and columns indicate the true distribution of times **t**. Colors indicate the number of features N. Features were chosen to be equal, specifically, we used 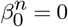 and 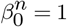. In both A and B, dashed lines correspond to the baseline where pseudotimes are known.

In Fig. 5A, we also illustrate the effect of the relationship between true pseudotime distribution **t** and priors *p*(*t*). The MSE of *β* is the closest to the regression baseline if the prior matches the true distribution of the data. However, if there is a disagreement, the MSE of *β* can even grow with *D*. One pessimistic interpretation is that the prior has a too sizable influence on the outcomes even for large donor sample size *D*. While this is true, at the same time, there is no way to get around this problem because of the inherent lack of identifiability when *N <* ∞. More optimistically, these results show that it is possible to encode our understanding of how donor sampling occurred if that knowledge is available.

### 5.2. Using multiple features

Fig. 5B shows that we can benefit from including informative features: as we increase *N*, the average MSE for each feature decreases, converging to the linear regression (known pseudotimes) baseline. For many features (e.g., *N* = 100), divergences from this baseline are observed only for many donors *D*, where MSEs are already low. This phenomenon can be understood in the light of our theoretical results: By virtue of Proposition 1, a higher number of features (with the same *β*) adding more features will lead to a more precise estimation of the (normalized) vector of pseudotimes *t*. These more precise estimates will, in turn, lead to estimates 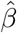 that are closer to the ones that would be obtained if pseudotimes were known.

In the previous experiment, we assumed that all features were equal. We also show that the same phenomenon still holds if features are not identical but are instead sampled from a distribution. To this end, we consider the scenario where 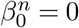 and 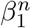 are sampled from a Gaussian distribution with mean *β*_1_ = 1 and variance *σ*^2^ ∈ *{*0, 10*}*. Results in Fig. 6 show that MSE for *β* is at a comparable scale independent of whether 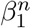 is random or not (*σ*^2^ = 0 v.s. *σ*^2^ = 10).

**Fig 6.**
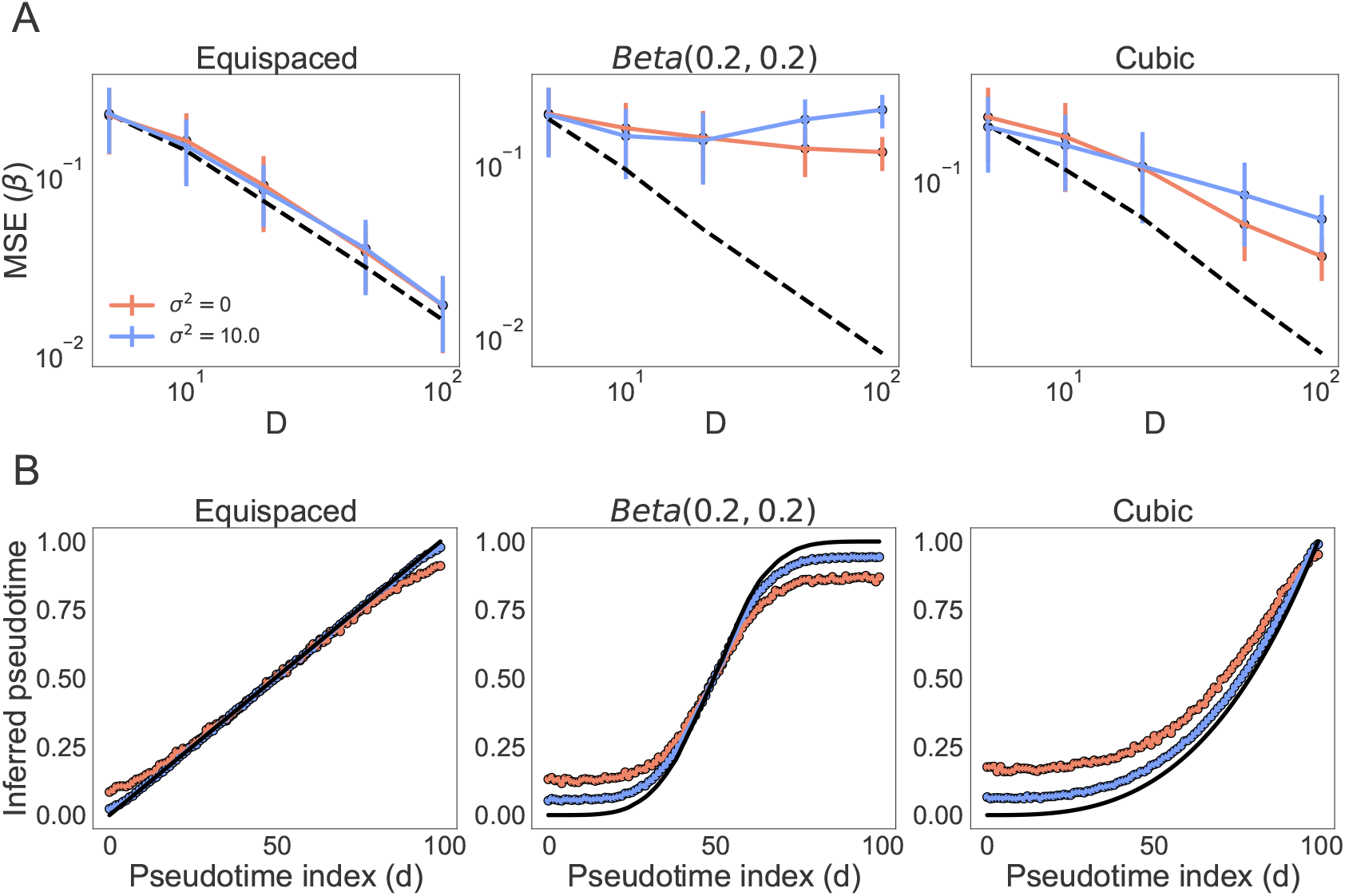
Effect of variability in β on estimation σ_2_ (colors) in linear model (Eq. (4)) with N = 10 features on parameter and pseudotime estimation. **(A) MSE of β = (β_0_, β_1_).** Dashed lines correspond to the baseline where the t_d_’s are known). (B) comparison of pseudotime estimates (averages over experiments)

However, unlike in the previous experiment, the added randomness in 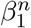 is beneficial for pseudotime estimation as we observe that time estimates in the *σ*^2^ = 10 case are consistently closer to true values. This may seem paradoxical as the *σ*^2^ = 10 corresponds to a noisier regime, and we may reasonably expect noisier results. Again, we can understand this phenomenon in the light of Proposition 1: larger values of *σ*^2^ translate into larger values of ||*β*||^2^, which in turn lead to a lower upper bound on the discrepancy between true and estimated pseudotimes, up to a normalization constant. This experiment highlights the relevance of feature diversity (i.e., variability in *β*) in tasks where pseudo time is the primary inferential goal. We will come back to this point when we discuss non-linearities.

### 5.3. Shrinkage in Gaussian and non-Gaussian models with hierarchical priors

While previous experiments address the multiple features case and the effect of variability in feature coefficients *β* on our estimates, none of the Bayesian models in those experiments explicitly model the sampling structure in coefficients. One natural question is whether the James-Stein phenomenon would manifest here, i.e., whether the MSE of *β* or *t* can be lowered using shrinkage estimators. These estimators, in turn, are obtained by imposing a suitable hierarchical structure on *β*. This question is not obvious since it depends on the interplay of *β* and the “nuisance” parameter *t*. In this experiment, we not only consider the Gaussian-linear model but include the more realistic non-linear negative binomial model with logarithmic link function (Eq. (2)).

As in the previous experiment, we assume that *β* are sampled i.i.d. from a superpopulation; i.e. 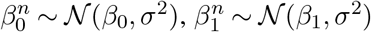). We consider Bayesian estimators based on four models: i) the one from previous experiments; i.e. a fully factorized independence prior on *β* (Fig 4A), and two hierarchical models (similar, but simpler than Fig 4B) where each 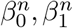 are sampled from a Gaussian distribution centered at *β*_0_, *β*_1_ and with variance 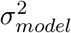, where 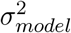 is either assumed known and equal to the true variance *σ*^2^ or treated as an additional parameter for which we perform Bayesian inference, using a half-normal hyperprior.

Fig. 7 shows that whether we benefit from a hierarchical model depends on the link function and the target parameter. As shown by Fig. 7B,C the hierarchical model is beneficial for the estimation of *β* in both the Gaussian and negative binomial case, although benefits tend to be more significant in the negative binomial model. More interestingly, regarding pseudotime estimation, benefits of shrinkage are observed only for the negative binomial model (Fig. 7A,C).

**Fig 7.**
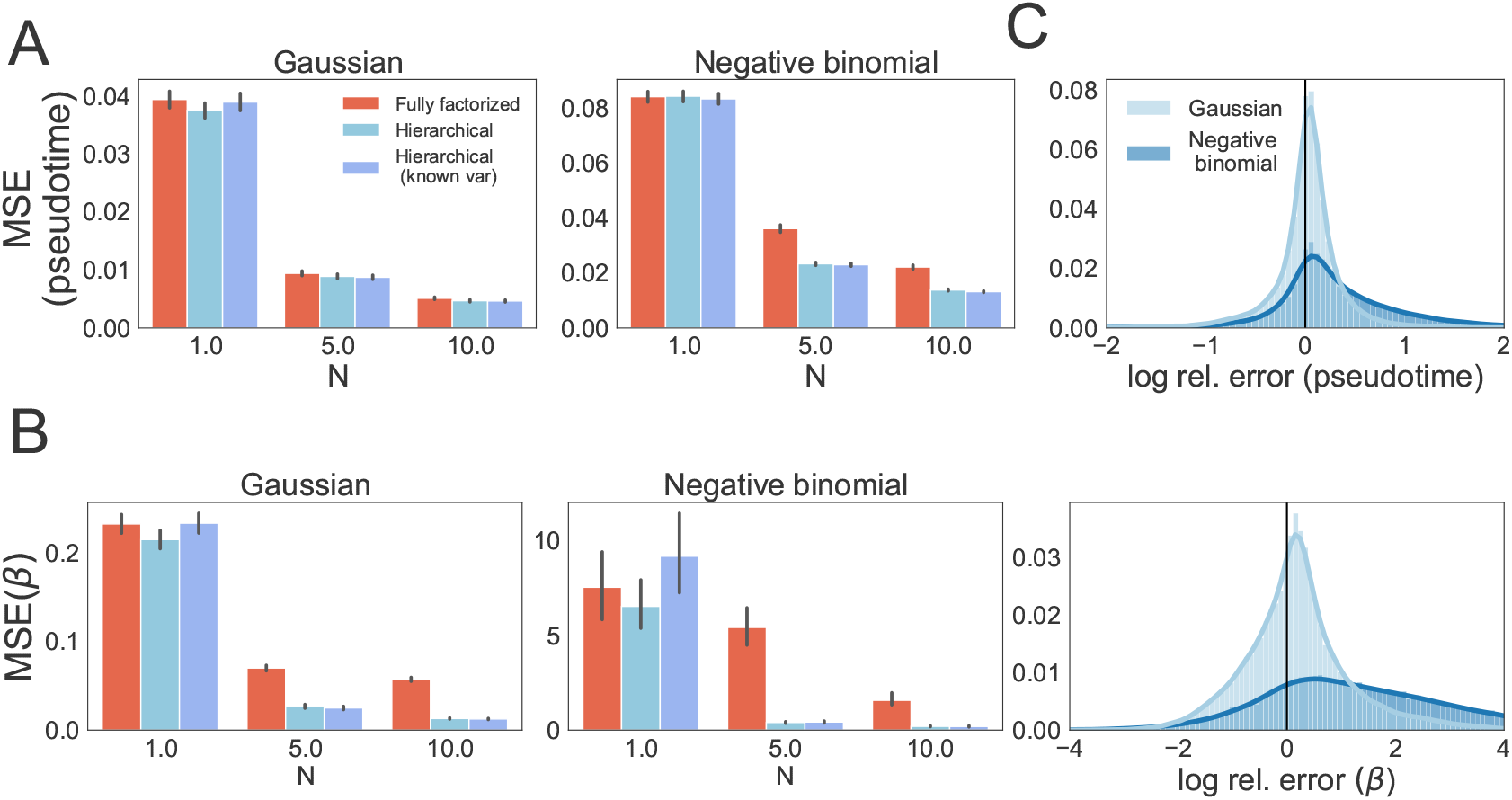
Estimation error of different hierarchical models for the linear-Gaussian model (Eq. (4)) and the negative binomial model (Eq. (2)). (A) MSE in pseudotime estimation. (B) MSE for β estimation. (C) Histograms of relative error (log-scale) for the fully factorized model compared to the hierarchical model each sample is an experiment). Each sample is an experiment

These results suggest that our framework can benefit from shared structure in *β* in two ways. First, through the “classical” James-Stein phenomenon whereby MSE on *β* is decreased by a hierarchical model on *β*. Second, at least in the negative binomial case, observe a less obvious “crossed’ shrinkage effect whereby estimation of pseudotime is benefited by the structure that we imposed on *β*. This may, in turn, explain a more significant benefit in *β* estimation in this case.

Additionally, Figs. 7A, B show that regardless of the observation model (Gaussian or negative binomial) and the fitted model (hierarchical or fully factorized), our estimators produce lower pseudotime MSE (average over different *t*_*d*_’s) and *β* MSE for larger values of *N*. Also, Supplementary Fig. S9 shows that a larger number of donors *D* has little impact on the average pseudotime MSE, but does help decreasing the *β* MSE. Moreover, Supplementary Fig. S9 shows that benefits of the hierarchical model in pseudotime estimation for the negative binomial model are constrained to a sufficiently small number of donors compared to the number of features. Additional results are shown in Supplementary Figs. S7 and S8.

### 5.4. Combining features with stage-specific information in non-linear models

One main feature distinguishing linear and non-linear models is that the signal-to-noise ratio SNR depends nonlinearly on times, implying that individual features can be preferentially informative about particular levels of AD pathology severity. Indeed, if we define (see Hastie et al. (2009))

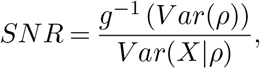

where *ρ* is the linear response (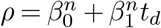 in Eq. (2)) and *g* is the link function (logarithmic in Eq. (2)), we observe that if 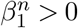, the SNR can be much larger at high compared to low severity levels, with an opposite behavior if 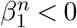.

We illustrate this phenomenon with an example of a negative binomial model with two features and an overdispersion parameter *A* = 100. The first feature has parameters 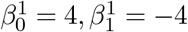, so the mean counts 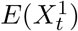 are large at low severity levels but decrease to eventually stabilize at high severity (see Supplementary Fig. S10 for examples of observations). The second feature has 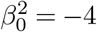 and 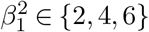 so the behavior is inverted, and the overall strength of this feature can be smaller 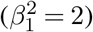, equal 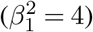 or larger 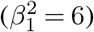 than the first feature.

We compare estimators based on each individual feature with the one based on both. Results are shown in Fig. 8. We observe that including both features leads to a lower MSE (on *t* and *β*) than any individual feature (Fig. 8A), suggesting that that the cross-talk of information between different features is beneficial for the estimation of each of them as they are uniquely informative about different aspects of the disease progression.

**Fig 8.**
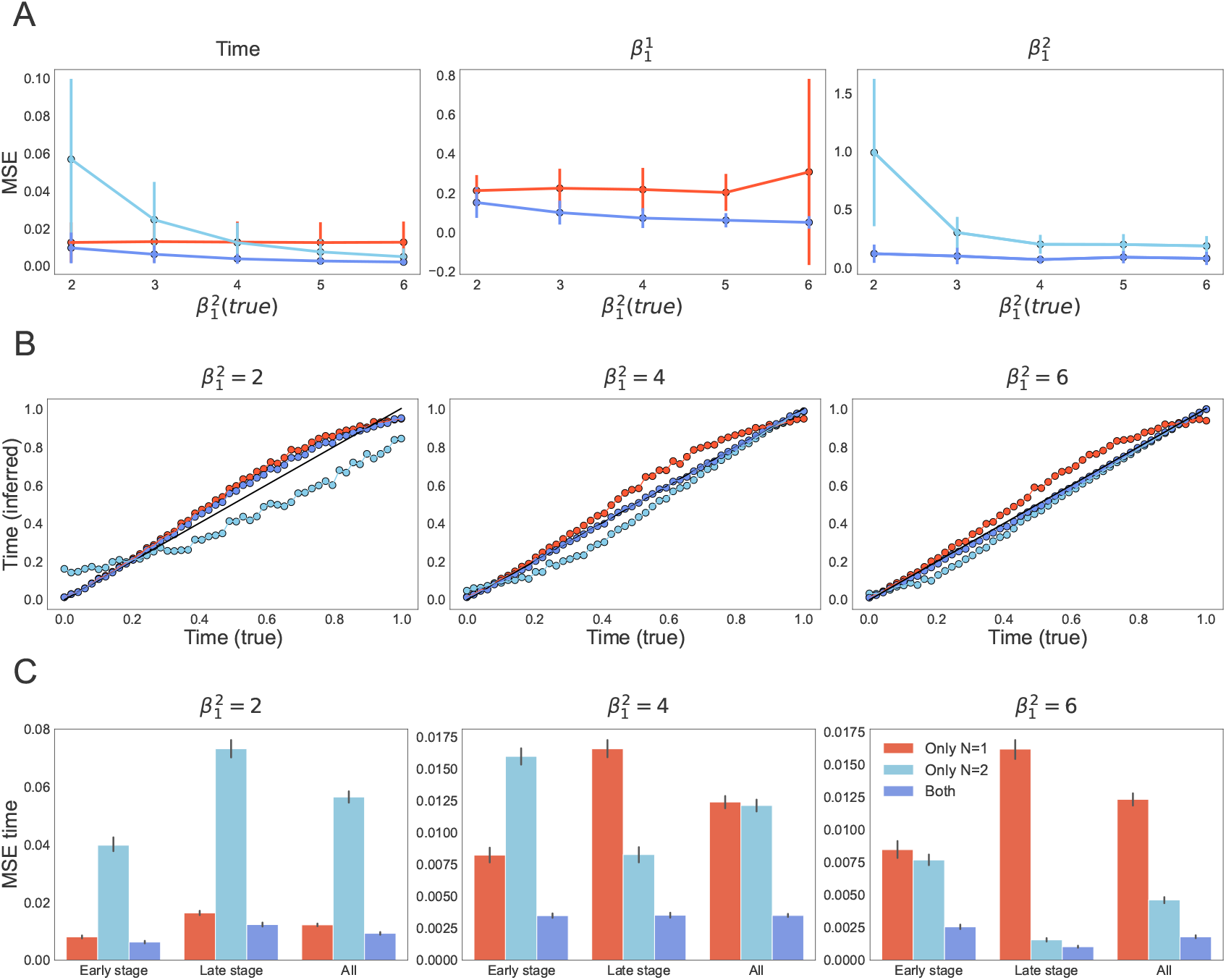
Inferring multiple features in the negative binomial non-linear model (Eq. (2)) (A): Average MSE for pseudotimes, and MSE for the strength of the first and second features (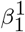 and 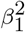) as a function of the strength of the second feature 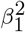 in models that only use individual features as well as the one that uses both (different colors). (B) Comparison of true and inferred pseudotimes (average over experiments) for different strengths of the second feature 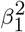. (C) Average MSE of inferred pseudotimes for the early and late stages, as well as their combination, for different strengths of the second feature 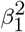.

We can better understand this phenomenon by looking at pseudotime sequences inferred by each model. As shown in Fig. 8B, the model that only uses the first feature accurately infers lower levels of severity but struggles in later-pseudotime inference, and the converse is true for the model that only uses the second feature. In contrast, the model that combines both accurately infers pseudotimes along the entire range. Fig. 8B also shows that this consensus will be achieved even if there is asymmetry (one feature stronger than the other) as long as there is enough signal in the system (i.e., if both 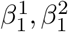 are sufficiently large). These observations are also supported by Fig. 8C, where we report MSE on pseudotime inference by classifying each donor as being on an ‘early’ (*t* ≤ 0.5) or ‘late’ (*t* ≥ 0.5) stage. Models with a single feature have the lowest MSE at the stages that the feature is informative about, and the overall pseudotime MSE of models containing the second feature decreases with the strength of this feature.

### 5.5. Additional experiments

In the Appendix (see Section “Supplementary experiments and experimental details”, we present more experiments showing that the phenomena described in linear models (Sections 5.1, 5.2, 4) also hold if we use Negative Binominal like-lihood (Eq. (2)). Additionally, we present supplementary experiments to illustrate the performance of the rate-optimal SVD-based estimators described in Sections 4 (Supplementary Figs. S1 and S2),as an alternative to the Bayesian estimators whose performance was studied in this section. In this context, we also show that the benefit of shrinkage estimators can be observed beyond the situations described here (see Supplementary Fig. S3).

## 6. Inferring latent pseudotime in the SEA-AD cohort

Finally, we apply our generative models (Fig. 4) to the problem of describing the progression of pathology aggregation in AD, and infer latent pseudotime for each donor in the SEA-AD cohort. As a reminder, our dataset consists of *D* = 84 donors in which *M* = 12 pathological markers are measured in each of *L* = 5 cortical layers of the MTG (as detailed in Section 2). After sampling from the models’ posteriors (see Section 3.3), posterior pseudotime estimates, *t*_*d*_, span the entire progression of disease (Fig. 9A), with an increased representation of later times, in accordance with the cohort distribution that is biased towards older ages and higher disease metrics (such as Braak, or ADNC, see Fig. 1B,C). Growth-rate parameter *β*_1_ and the parameter quantifying the initial seed levels of pathology *β*_0_, revealed that canonical AD protein markers follow increasing exponential dynamics from small values, while, as expected, a marker quantifying the number of neuronal cells per unit area decreases as AD progresses (Fig. 9B,C).

**Fig 9.**
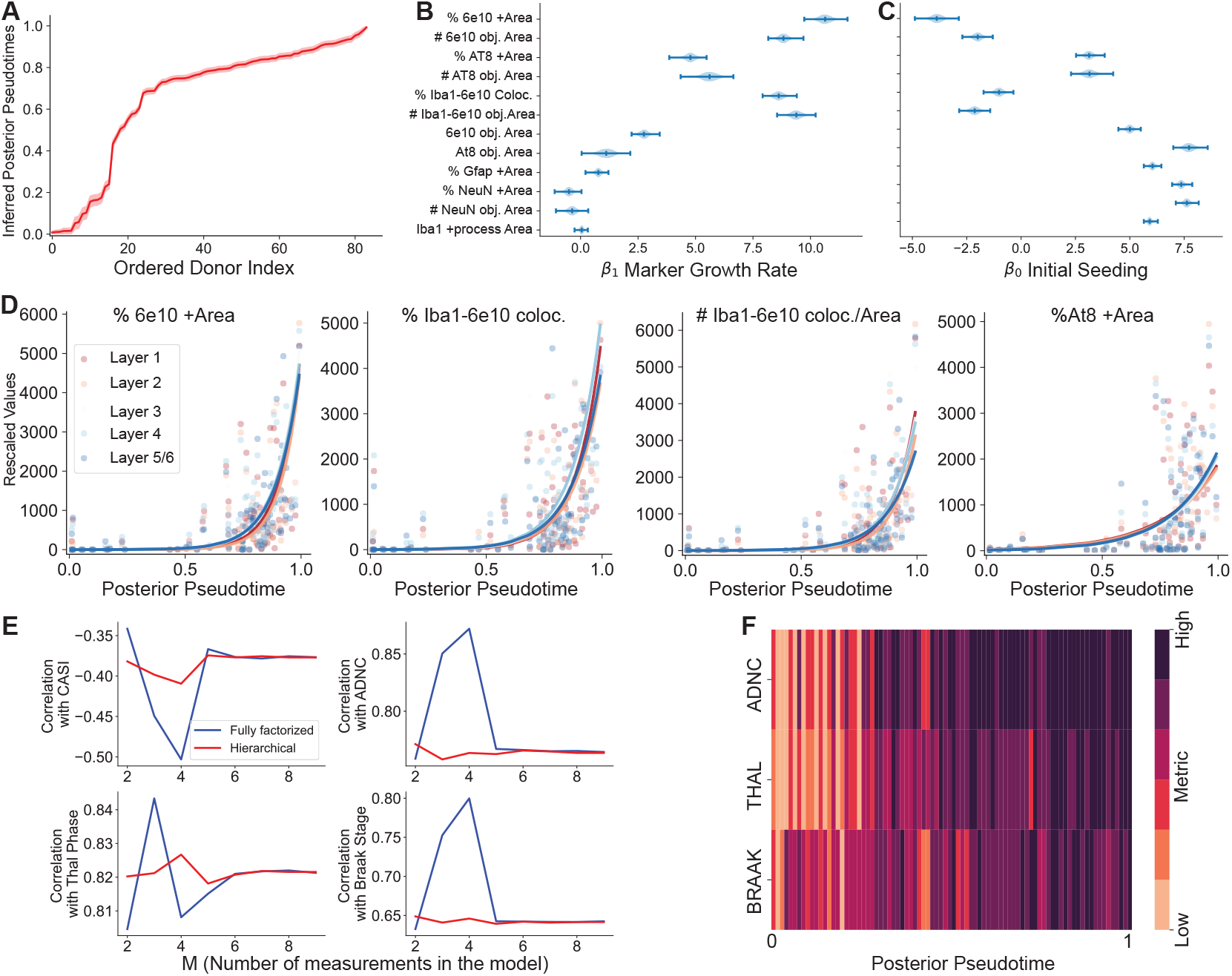
Posterior distributions of model parameters for SEAD-AD data set of pathological proteins. (A) Inferred pseudotimes t_d_ for donors sorted in ascending order. B) posterior distribution of β_1_, growth-rate and (C) β_0_, initial seeding model parameters. In each case, error bars denote the distribution of posterior means inferred for each layer. (D) Posterior means (solid lines) plotted in conjunction with the data for 4 exemplary pathological markers. Color-codes indicate observations for each cortical layer. **Inferred pseudoprogression correlates with existing metrics for pathology progression**. (E) Spearman correlation between the associated progression metric and inferred pseudoprogression score td, as the number of measurements to infer td increase. Both models (fully factorized and hierarchical) show high correlations with these metrics. However only a few measurements are required to maximize the correlations. (F) All relevant metrics increase monotonically with the inferred pseudo-progression score inferred from our model

Consistent with our predictions from simulation studies, we observed that for *T* = 84 donors, both the hierarchical model and the fully-factorized model result in similar posterior distributions of model parameters. This is due to the size of the dataset in which a sufficient number of donors compensate for the extra information provided by the layer and measurement covariance incorporated by the hierarchical model. Quantitative metrics comparing both models include the expected log posterior predictive distribution (Vehtari, Gelman and Gabry, 2017) and the Watanabe Information Criteria (WAIC, Watanabe and Opper (2010)). These metrics indicate that the hierarchical model better fits the data but the gain at this data regime is marginal (Supplementary Figures S11 and S12).

Next, we evaluated our model through correlations of our posterior pseudotime estimates with established neuropathological assessments, gaining the ability to independently validate our estimates with orthogonal metadata not used to fit the model. We observe that the pseudotime parameter is highly correlated with neuropathological assessments Braak, Thal, and ADNC (positive correlation) and cognitive scores CASI (negative correlation, Teng et al. (1994b)) (Fig. 9 E,F), These orthogonal metrics describe the brain-wide progression of multiple pathological proteins or quantify cognitive decline (see Section 2).

The definition of the Braak stages asserts that Braak stage IV (or higher) is identified by the presence of pTau pathology in the MTG, as described by Braak et al. (2006). This relationship aligns our inferred pseudotime, which is based on pTau measurements, with the Braak staging system. Nevertheless, this correlation is only moderate (the Spearman correlation between the inferred pseudotime and a binarized Braak score—0 for stages below 4 and 1 for stages 4 and above—is 0.35) when compared to the correlation using all Braak stages (correlation >0.65). This suggests that pseudotime is effectively ordering donors across the entire range of Braak stages.

### 6.1. Exploring the importance of pathological markers as model regressors

We next investigated the contribution of each feature to our latent pseudotime. As our problem does not map to a regression problem, in which variable importance has been extensively studied, we resorted to multiple metrics to quantify each pathological marker contribution. To compare the effect of each pathological marker as an explanatory variable, making an analogy to regression problems, we first look at the growth-rate parameter *β*_1_. Parameter values differ considerably across all *M* = 12 features, with six having the highest absolute value (Fig. 10A). However, in our case, parameter magnitude could be dependent on the signal-to-noise ratio in each feature and be loosely related to pseudotime inference. To find a different way of quantifying associations between pseudotime and each measurement, we utilized calculation of the mutual information (MI), understanding MI as a means to identify feature importance in nonlinear models (Beraha et al., 2019). MI identifies the 7 features as the most relevant ones to inform pseudotime inference (Fig. 10B).

**Fig 10.**
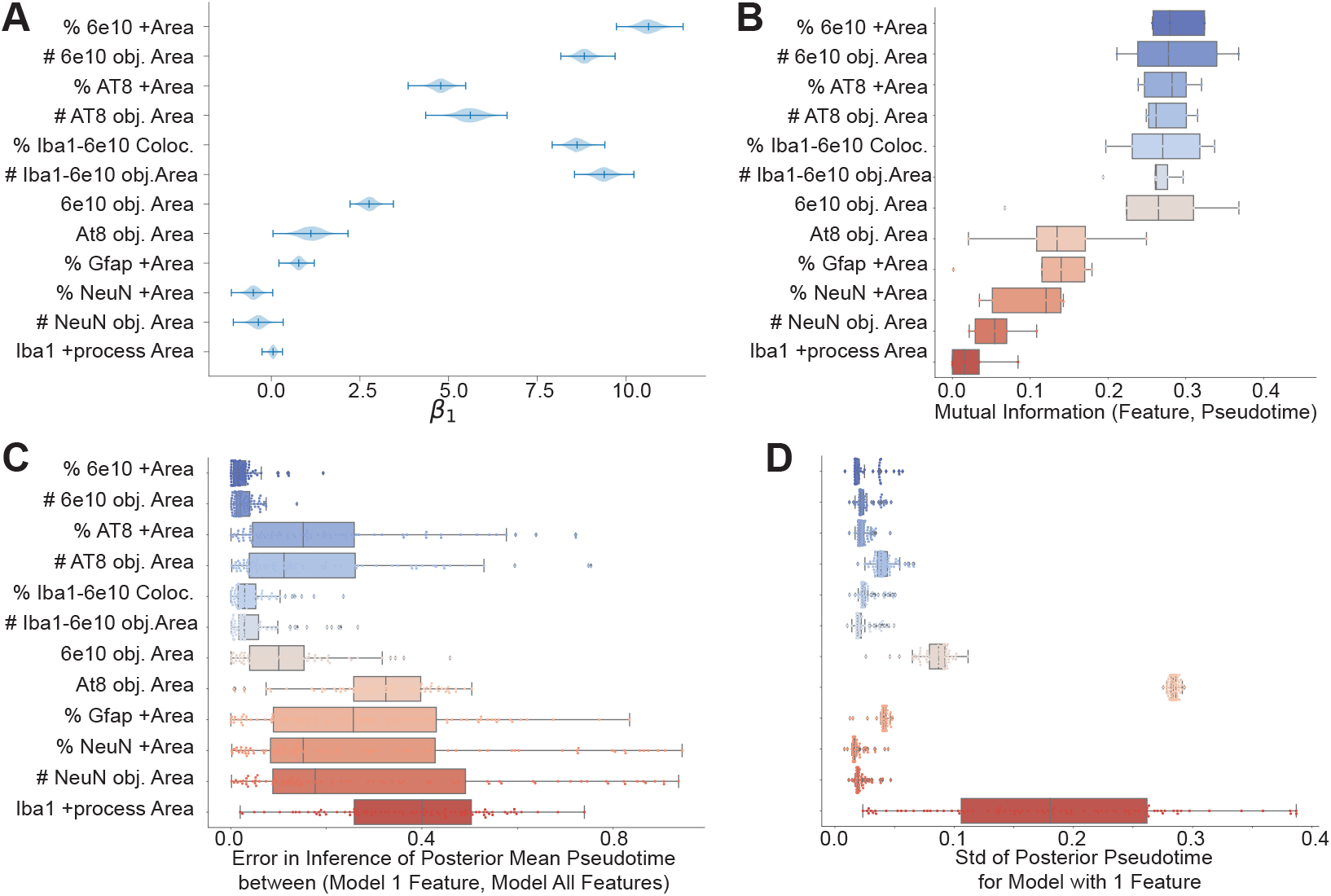
Multiple metrics describing the importance of each pathological feature to inform latent pseudotime inference. (A) Growth-rate parameter, β_1_, for each pathological feature, ordered according to decreasing mutual information (MI). (B) MI between each pathological feature and posterior pseudotime. (C) Error in pseudotime inference by comparing posterior times inferred using one feature vs all M = 12 features. (D) The standard deviation of posterior estimate, t_d_, for each single-feature model.

As an additional feature importance computational experiment, we calculated pseudotime posterior estimates for models using each feature in isolation. Then, we calculated the error in the posterior pseudotime parameter *t*_*d*_ inferred from only one feature compared to all fea-tures, (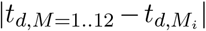 (Fig. 10 C) and the standard deviation in the posterior distribution of each pseudotime parameter *t*_*d*_, to show posterior uncertainty on pseudotime estimation (Fig. 10D). Our previous simulation studies informed us that features with high growth-rate parameter *β*_1_ would have a high-dynamic range of pathology and observations with high pathological values. This in turn is more informative for obtaining identifiable estimates of *t*_*d*_. Our computational results are in accordance with our simulation studies, revealing that the first seven pathological variables (order according to decreasing MI) have small pseudotime error and variance (Fig. 10C-D). Taken together all these computational experiments suggest that the seven most informative variables are a more compact subset able to create reliable pseudotime posterior estimates.

### 6.2 Informing Future Experimental Design

This section is driven by the aim of leveraging our models to guide the design of future experiments. While this aim could be framed as a Bayesian optimization problem, we opt for a qualitative approach guided by several principles. Firstly, the process of collecting, processing, and analyzing neuropathological measurements, as presented here, is labor-intensive. Our objective is to utilize the insights gained from the preceding section to identify a concise set of highly informative variables. Although reducing the feature set could be in detriment of the quality of fit (as described in our Simulated Studies Section 5), this will streamline the time required for future experiments. Secondly, the selection criteria for brain donors in the SEA-AD cohort resulted in a bias towards older donors and those at advanced pathological stages. Most donors are of old age, mean age = 88 years old, and exhibit an inferred pseudotime greater than 0.5. In the future, we would like to diversify the SEA-AD cohort by including donors that uniformly span the entire pseudotime spectrum (from 0 to 1). A balanced cohort, representative of all disease stages, facilitates downstream analysis of multiple disease epochs. Moreover, pseudotime being an inferred parameter poses challenges in its use for donor preselection and cohort balancing. To address this, we aim to enhance the model by incorporating other observable variables such as donor metadata or staging information. This approach will enable us to strategically sample donors and mitigate biases.

To achieve these goals, we investigate a model incorporating *M* = 7 pathological features, chosen for their high standardized beta coefficient, mutual information, and low error and standard deviation in time inference (see Fig. 10). These features constitute a minimal set as desired. To suggest future brain donors based on easily measurable variables, we augment the model with categorical staging metrics (Braak, Thal, and ADNC). Let *m* represent a staging metric, *d* denote a donor, and *k* signify a one-hot encoded stage level, then the vector 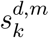 represents:

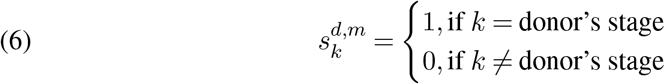

For example, Thal phases range from phase 0 to 5, with increasing phases indicating an increasing number of brain regions containing amyloid-beta pathological burden. A donor exhibiting phase-2 Thal phase would be represented as *s*^*d,T hal*^ = [0, 0, 1, 0, 0, 0]. We incorporate staging metrics into our formalism and model them as:

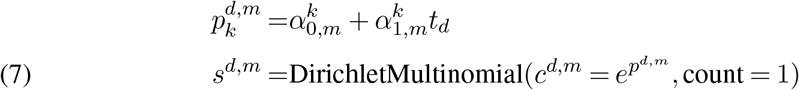

Assuming that pseudotime *t*_*d*_ = 0 aligns with all staging metrics being zero and that their values are monotonically higher, we assign prior parameters *α*_0_ and *α*_1_ for each stage *k*.

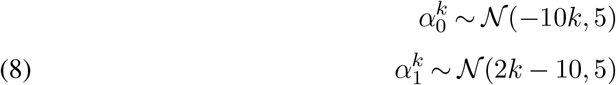

Next, we perform Bayesian posterior inference and compare posterior estimates of the augmented model (Augmented, *M* = 7 + Staging metrics) against our model that includes all features (All Feats, *M* = 12) in Fig. 11 and Supplementary Fig. S15. Both models resulted in similar posterior estimates, revealed by comparing posterior pseudotime estimates for each donor and dynamic parameters (*β*_0_, *β*_1_) across measurements common to both settings (Fig. 11A-B). In both cases, neuropathological data used to infer posterior parameters results in concordant estimates and relatively small posterior standard deviations. We observed that staging information does not significantly impact pseudotime posterior estimates, both estimates concentrating around the same mean, when most informative neuropathological features are selected. Although speculative, this may be indicative of the fact that the staging information may be too coarse to provide any additional information on disease progression on top of the one already provided by neuropathological markers. At the same time, the augmented model is able to effectively model staging metrics, observed when comparing staging metrics’ posterior predictive distributions and observed data (Fig. 11C). Taken together, these results highlights the ability of our augmented model together with the set of informative features to accurately model pseudotime and AD staging metrics in concert.

**Fig 11.**
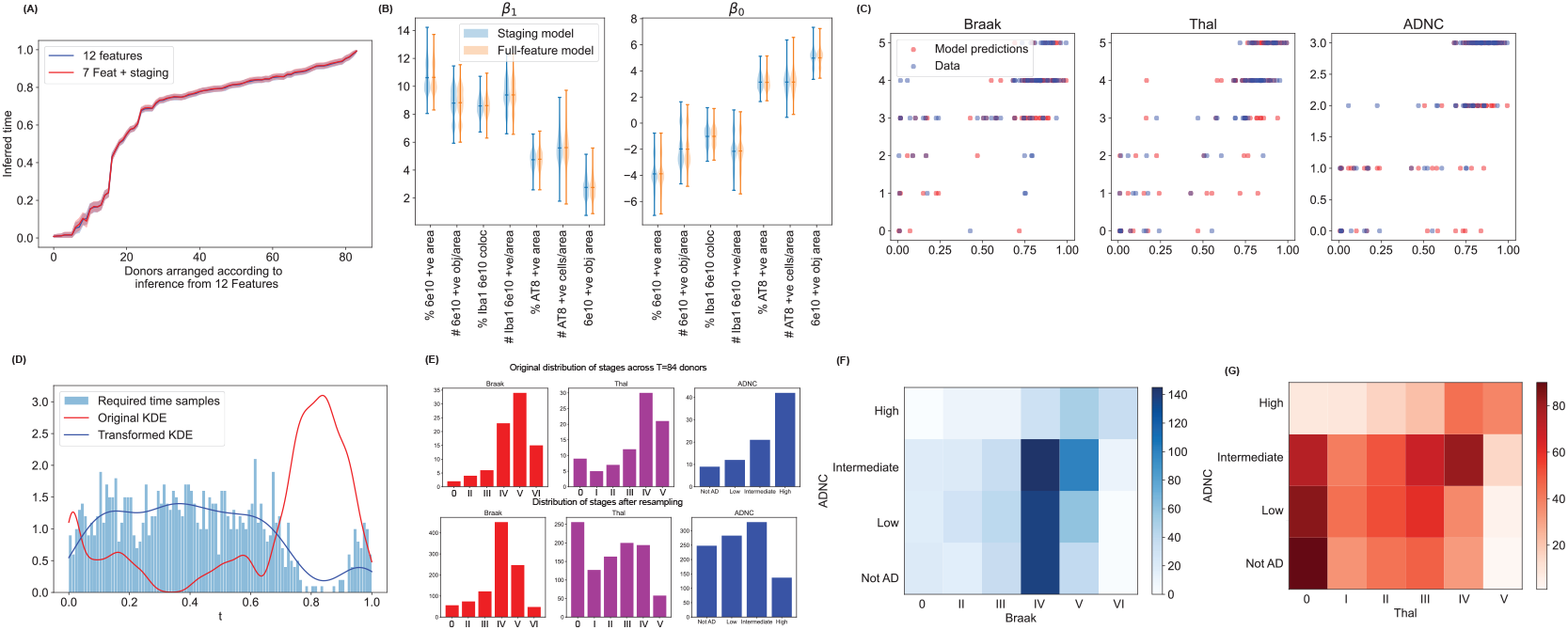
Model augmented with staging metrics is able to effectively represent experimental data. (A) Comparison of posterior time estimates predicted by models spanning all features and features plus staging metrics. (B) Posterior β_0_ and β_1_ distributions (concatenating values from all layers and samples) in both models (C) Posterior predictive estimates reveal effective modeling of staging metric (Braak, Thal and ADNC). **Augmented Model incorporating staging information informs experimental design to balance SEA-AD cohort across disease stages** (D) To balance SEA-AD cohort, we used kernel density estimates to describe the distribution of inferred pseudotime values. Next, we empirically calculate a pdf corresponding to the least probable time values. N = 1000 samples are drawn from this distribution and the resulting histogram is displayed (E) Pseudotime samples from (D) are then converted to the associated Braak, Thal and ADNC stage. Top plot shows the distribution of stages in the donor dataset, bottom plot shows the distribution of stages corresponding to the resampled times. (F,G) Heatmaps showing the Braak/ADNC and Thal/ADNC joint histograms. The associated distribution of stages can aid experimentalists in improving donor sampling to balance their cohort.

Finally, we utilize our augmented model to guide experimental design, aiming to achieve balance (equal representation of donors across pseudotime) in the SEA-AD cohort. To accomplish this, we first estimate the distribution of posterior pseudotimes, *p*(*t*), by 1) aggregating mean posterior pseudotime estimates for each donor, 2) constructing a histogram, and then 3) applying kernel density estimation. Next, we approximate the unnormalized distribution of under-sampled pseudotimes in the SEA-AD cohort by computing *p*_*under*_(*t*) = 1 − *p*(*t*). This approximation enables us to generate samples of under-sampled pseudotimes (*n*_*samples*_ = 1000, see Fig. 11 D). These values can then be employed to compute posterior predictive staging metrics based on the posterior estimates of staging parameters (see Fig. 11E-G). These predictions inform the staging metrics of suggested new donors that should be included in the study to achieve cohort balance. Given the current distribution of our cohort, it is apparent that sampling from early and middle stages would be advantageous.

## 7. Discussion

We introduced a Bayesian framework to model Alzheimer’s Disease progression, incorporating a latent pseudotime variable to depict disease evolution. We modeled observations utilizing biophysically-inspired assumptions that posit pathological aggregation follows exponential dynamics. Our inference method is useful for datasets restricted to cross-sectional data, consolidating observations and inferring a single and common disease trajectory. We demonstrated the utility of our framework using simulation studies and real datasets originating from the SEA-AD consortium, profiling AD neuropathological proteins in a brain region. Applying our model to the SEA-AD cohort dataset, consisting of 84 donors, allowed us to infer donor pseudotime and pathological protein dynamics. Pseudotime estimates are biased towards late values, in accordance with our observations that the SEA-AD cohort is biased towards older donors and high pathology. Posterior predictive checks validate the goodness of our fits. In addition, comparisons with existing neuropathological staging-based metrics reveal that pseudotime ordering aligns well with brain-wide neuropathological assessments, indicating that our model is able to map AD trajectory when detailed information from only one brain region is used (the Middle Temporal Gyrus).

Next, we assembled a collection of metrics to assess the impact of each pathological protein on pseudotime inference, identifying the most informative features. These features exhibited high growth rates, mutual information between their values and pseudotime, and individually exhibited precise predictions of pseudotime with low prediction error. Incorporating these highly informative variables alongside neuropathological assessments, we enhanced our model to deduce the latter. When integrated with staging data, our enhanced model can guide the identification of neuropathological stages for future donors to be included in the existing SEA-AD cohort, ensuring a balanced representation of donors across all pseudotimes sand enhancing downstream analysis across the disease spectrum. While our work assumes that the pathology markers align donors along the same pseudotime trajectory, which is true for the MTG region of the brain, this assumption may not hold true for other brain regions. This would require approaches that can overcome low-data regimes to successfully leverage all regions in a combined overarching multi-regional pseudotime.

Our results demonstrated the success of our methodology, supported by our theoretical explorations. In our work, we derived theoretical results on a simplified linear model to understand the best possible performance. A stimulating recent line of work has provided algorithmic (Kidzinski et al., 2022) and statistical (Chen, Li and Zhang, 2020; Du, Wasserman and Roeder, 2023) guarantees for matrix factorization problems in generalized-linear setups, a setup akin to ours. However, these results by themselves are not sufficient to explain the richness of phenomena manifesting in our setup. Future studies should study the simplified model and extend Proposition 2 by deriving explicit rate bounds for the signal strength estimates, and also extend our theoretical results to a generalized linear model case. We made contributions in this direction through extensive simulations and showed that whether or not we benefit from hierarchical models for pseudotime estimation depends on the type of model and error distribution, and benefits are only observed beyond linear models with Gaussian errors. Future studies should develop a theoretical framework to explain this phenomenon.

## Acknowledgments

The authors would like to thank Sivaraman Balakrishnan for useful theoretical discussions, Bob Carpenter for insightful discussion on modeling and inference approaches, Iris Stone for manuscript feedback, and the entire SEA-AD team for creating an open source resource for the study of Alzheimer’s Disease.

