## Supplementary Material for "B-BIND: BIOPHYSICAL BAYESIAN INFERENCE FOR NEURODEGENERATIVE DYNAMICS"

*Submitted to the Annals of Applied Statistics*

### APPENDIX A: DETAILED DERIVATION OF BIOPHYSICAL MODEL OF PRIONIC AGGREGATION AND PROPAGATION

In addition to local prionic aggregation and fragmentation, the toxic seeds spread to other connected brain regions. For analyzing a spatially-limited region in the brain e.g. a single anatomical region such as the MTG, it can be shown that the time scales of seed spreading are slower than the local seed multiplication through aggregation-fragmentation dynamics (Meisl et al. (2021)). In this limit, we can consider an *in vitro* model with spatial gradients ignored. With an *in vitro* model, we can add additional specifications to separately analyze the time evolution of prionic number and prionic size.

The kinetic equations that describe the evolution of the monomeric unit number (or concentration) $m_0$, and the evolution of toxic prionic aggregates with $i$ units $y_i$ can be written as (Masel, Jansen and Nowak (1999)):

(S1a) $$\frac{dm_0}{dt} = -\kappa m_0 y + 2b \sum_{i=1}^{n-1} \sum_{j=i+1}^{\infty} i y_j$$

(S1b) $$\frac{dy_i}{dt} = -\kappa m_0 (y_{i-1} - y_i) - b(i-1)y + 2b \sum_{j=i+1}^{\infty} i y_j$$

where $b$ is the fragmentation rate per length of aggregate and $\kappa$ is the monomer association rate (polymerization rate). $y = \Sigma y_i$ is the total prionic aggregate concentration, and $n$ is the size of stable prionic aggregates, so that any aggregate below this critical size is unstable and rapidly converts to monomeric units $m_0$.

To obtain the aggregate number/concentration as a function of time ($y = \Sigma y_i$), we close the equations above using summation and first moment $z = \Sigma i y_i$ to get differential equations in $m_0, y, z$:

(S2a) $$\frac{dm_0}{dt} = -\kappa m_0 y + b(n)(n-1)y$$

(S2b) $$\frac{dy}{dt} = bz + b(2n-1)y$$

(S2c) $$\frac{dz}{dt} = \kappa m_0 y - b(n)(n-1)y$$

Usually, it is assumed that the total mass of monomeric units available ($M_{tot} = m_0 + z$) is constant over time, i.e. there is no production nor degradation of monomeric/polymeric units. To obtain the general behavior for $x(t), y(t), z(t)$ and $s(t)$, we solve these equations for early-time and late-time dynamics.

The initial time solution for y (concentration or number of prionic aggregates) can be expressed as (Masel, Jansen and Nowak (1999)):

(S3) $$y \sim y(0)e^{\beta t}$$

where $\beta \sim \sqrt{\kappa M_{tot} b}$ is the rate constant that depends on the fragmentation-based seeding rate $b$, total (prionic and healthy) pool of protein monomer concentration $M_{tot}$ and polymerization rate $\beta$. In addition, the time evolution of the size of misfolded aggregates, $s = z/y$, can be derived from equations for $z$ and $y$. For initial time ($m_0 \sim M_{tot}$), the size evolution is:

(S4) $$\frac{ds}{dt} = \kappa M_{tot} + b(s)(1-s)$$

which follows sigmoidal dynamics.

For later times, we can instead assume $M_{tot} \sim z$ to obtain

$$\frac{dy}{dt} = b(M_{tot} - y) \tag{S5}$$

to get $y \sim M_{tot} - e^{-bt}$. If we assume that the total aggregate concentration doesn't reach this saturation level for most intents and purposes, then the aggregate concentration can be modeled as increasing exponentially in time.

The simplest model that can incorporate the spatiotemporal dynamics of aggregate growth and spread can be expressed as a Fisher-Kolmogorov-Petrovsky-Piskunov equation (Meisl et al. (2021); Weickenmeier, Kuhl and Goriely (2018); Cohen et al. (2014)):

$$\frac{dc(r,t)}{dt} = D\nabla^2 c(r,t) + \beta c(r,t)(1 - c(r,t)) \tag{S6}$$

where $D$ is the effective diffusion constant, and $\beta$ is the effective multiplication rate (Meisl et al. (2021)) incorporating aggregation and fragmentation kinetics.

### APPENDIX B: PROOFS

Here we prove Propositions (1) and (2). We start with a preamble on the main mathematical definitions. We compare subspaces using the standard metric between orthogonal spaces $U_1$ and $U_2$ in $\mathbb{R}^{p\times r}$,

$$D_F(U_1, U_2) = \inf_{O \in O_r} \left\| U_1 - U_2 \right\|_F,$$

where $F$ is the Frobenious norm and $O_r$ is the set of Orthogonal matrices of dimension $r$. In the $r = 1$ case this metric reduces to

$$D_F(U_1, U_2) = \min\{\left\|U_1 - U_2\right\|_2, \left\|U_1 + U_2\right\|_2\}.$$

Our results are based on $\sin\Theta$ distances (Cai and Zhang, 2018). These are defined in the following way. For two orthogonal subspaces $U_1, U_2 \in \mathbb{R}^{p\times r}$, suppose the singular values of $U_1^T U_2$ are $\sigma 1 \geq \ldots \geq \sigma_r \geq 0$. The principal angles between $U_1$ and $U_2$, $\Theta(U_1, U_2)$, is the diagonal matrix:

$$\Theta(U_1, U_2) \equiv diag\left(\cos^{-1}(\sigma_1), \cos^{-1}(\sigma_2), \cdots, \cos^{-1}(\sigma_r)\right).$$

Based on these angles, we may define the $sin\Theta$ distance between subspaces, for example, through the quantity $||\sin\Theta(U_1, U_2)||_F$. By (Cai and Zhang, 2018, Lemma1) this $\sin\Theta$ distance is related to the above subspace distance $D_F$ through where

$$D_F(U_1, U_2) \leq \sqrt{2}\left\|\sin\Theta\left(U_1, U_2\right)\right\|_F = \left\|U_1 U_1^\top - U_2 U_2^\top\right\|_F. \tag{S7}$$

**B.1. Proof of Proposition 1.** To obtain estimation bounds for $t'$ and $\beta'$ we turn to specialized risk analysis for subspace estimation Cai and Zhang (2018) based on variants of the well-known Wedin's $\sin\Theta$ perturbation theorem Wedin (1972)

THEOREM B.1 (Theorems 3 and 4 in Cai and Zhang (2018)). *Suppose that we observe $X = Y + Z$ where $Y$ is of rank $r$ and $Z$ is a matrix of independent zero-mean Gaussian variables with variance $\phi^2$. Let $Y = U\Sigma V^\top \in \mathbb{R}^{D\times N}$ be the singular value decomposition (SVD) of $Y$. Likewise, $X = \hat{U}\hat{\Sigma}\hat{V}^\top$ be the SVD of $X$.*

*If $\sigma_r(Y)$ is the $r$th largest singular value of $Y$, there is an absolute constant $C$ such that*

$$E\|\sin\Theta\left(V,V^{\top}\right)\|_F^2 \le \frac{CN\phi^2\left(\sigma_r^2(Y)+D\phi^2\right)}{\sigma_r^4(Y)} \wedge r$$

$$E\|\sin\Theta\left(U,U^{\top}\right)\|_F^2 \le \frac{CD\phi^2\left(\sigma_r^2(Y)+N\phi^2\right)}{\sigma_r^4(Y)} \wedge r,$$

*Additionally, let $\mathcal{F}_{r,\delta}$ be the set of rank $r$ matrices $Y=U\Sigma V^{\top}$ with $\sigma_r(Y)\ge\delta\phi^2$. If $r\le D/16\wedge N/2$, then for some $c>0$*

$$\inf_{\tilde V}\sup_{X\in\mathcal{F}_{r,\delta}} E\|\sin\Theta\left(V,V^{\top}\right)\|_F^2 \ge \frac{c\phi^2 N\left(\delta^2+D\phi^2\right)}{\delta^4}\wedge r$$

$$\inf_{\tilde U}\sup_{X\in\mathcal{F}_{r,\delta}} E\|\sin\Theta\left(U,U^{\top}\right)\|_F^2 \ge \frac{cD\phi^2\left(\delta^2+N\phi^2\right)}{\delta^4}\wedge r.$$

Let's apply the above results with $r=1$, $U=t'$, $V=\beta'$ and $\Sigma=\lambda=||\beta||_2||t||_2$. Using the bound (S7) we obtain

$$ED_F(\hat\beta',\beta')^2 \le \frac{2C\left(||\beta||_2^2||t||_2^2+D\right)}{||\beta||_2^4||t||_2^4}\wedge 1$$

And, therefore, by a suitable division by $1/N$ and $1/D$ we obtain

$$\frac{1}{N}ED_F(\hat\beta',\beta')^2 \lesssim \frac{\left(\overline{\beta}_N^2\overline{t}_D^2+\phi^2\frac{1}{N}\right)}{(\overline{\beta}_N^2\overline{t}_D^2)^2}\wedge 1.$$

The bound with $t'$ is completely analog. By the same argument, we can also derive the lower bounds.

**B.2. Proof of Proposition 2.** The proof is divided into two steps. First, we show that we can identify the product $t_\infty\times\beta_\infty$, where

$$\beta_\infty^2\equiv\lim_{N\to\infty}\bar\beta_N^2,\quad \bar\beta_N^2,\equiv||\beta||^2/N,$$

and

$$t_\infty^2\equiv\lim_{D\to\infty}\bar t_D^2,\quad \bar t_D^2\equiv||t||^2/D.$$

Second, we show that $t_\infty$ is identifiable, from which the identifiability of $\beta_\infty$ follows trivially.

**Identification of $t_\infty\times\beta_\infty$**
Call $\lambda_{D,N}=\bar t_D^2\times\bar\beta_N^2$. We analyze separately two cases, $\phi^2$ is known or $\phi^2$ unknown. In the case where $\phi^2$ is known, we consider the following estimator of $\lambda_{D,N}$:

$$\hat\lambda_{D,N}=\max\left\{\frac{||X||_F^2}{DN}-\phi^2,0\right\},$$

where $||X||_F$ is the Frobenious norm of matrix $X$. We also have

$$||X||_F^2=DN\lambda_{D,N}+\langle\beta t^{\top},\epsilon\rangle_F+||\epsilon||_F^2.$$

The last term is a sum of $D\times N$ $\phi^2$-scaled chi-square random variables with one degree of freedom so that as $N,D$ grow to infinity,

$$\frac{||\epsilon||_F^2}{ND}\to\phi^2.$$

Additionally, by independence of $\epsilon$,

$$\frac{1}{ND}\langle \beta t^\top, \epsilon\rangle_F = \frac{1}{ND}\sum_{d,n}\beta_n t_d \epsilon_{n,d} \sim \mathcal{N}\left(0, \frac{1}{ND}\bar{\beta}_N^2 \bar{t}_D^2\right).$$

Therefore, as $\bar{\beta}_N^2$ and $\bar{t}_D^2$ are bounded (these sequences have limits), we conclude that this inner product converges almost surely to zero. Putting it all together,

$$\lim_{D,N\to\infty} \hat{\lambda}_{D,N} = \lim_{D,N\to\infty} \lambda_{D,N} = t_\infty \times \beta_\infty.$$

If $\phi^2$ is unknown, it suffices to show that it can be consistently estimated from data. Then, we can modify our estimator $\hat{\lambda}_{N,D}$ by replacing $\phi$ by a consistent estimator of $\hat{\phi}$. Consistent estimation of $\phi^2$ in our setup has already been studied in (Kritchman and Nadler, 2009) if $D/N \to c \in (0,1)$. These estimators rely on the analysis of the spectrum of $XX^T$. By contrasting the largest and remaining eigenvalues of $XX^T$ it is possible to separate the signal component $t_\infty \times \beta_\infty$ from noise.

**Identification of $t_\infty$, $\beta_\infty$**

We can identify $t_\infty$ from the constraint that the support of $t$ is the interval $[0,1]$. We will employ a limit argument where we need to consider a limit of true sampled sequences $t$ and $\beta$ as $N$ and $D$ grow to infinite. In practice, this is only a technical detail and we will avoid unnecessary double indexing of sequences.

By Proposition 1, as the number of features $N$ goes to infinity, for each number of donors $D$, the vector of normalized estimated pseudotimes $\hat{t}$ converges almost surely to the normalized sequence of times $t'$ (up to a sign). Therefore, in the limit we can make the assignment $\hat{t} = st'$ for $s \in \{-1, 1\}$. relate to $t'$ by $t = Kt'$ for a certain constant $K \neq 0$. The support constraint implies that almost for any chosen sequence of times $t_d$

$$\lim_{D\to\infty} \max_{d\in[1,D]} t_d = 1 \quad \text{and} \quad \lim_{D\to\infty} \min_{d\in[1,D]} t_d = 0.$$

Therefore, the estimator

$$\hat{\hat{t}} = \frac{|\hat{t}|}{\max_{d\in[1,D]} |\hat{t}_d|}$$

must equal $t$ as $D \to \infty$ since

$$\begin{aligned}
\lim_{D\to\infty} \hat{\hat{t}} &= \lim_{D\to\infty} \frac{|\hat{t}|}{\max_{d\in[1,D]} \hat{t}_d} \\
&= \lim_{D\to\infty} \frac{|t'|}{\max_{d\in[1,D]} |t'_d|} \\
&= \lim_{D\to\infty} \frac{|t||K^{-1}|}{|K^{-1}| \max_{d\in[1,D]} |t_d|} \\
&= t \frac{1}{\lim_{D\to\infty} \max_{d\in[1,D]} t_d} \\
&= t.
\end{aligned}$$

We have shown that $t$ (and not only $t'$) can be identified. Therefore, the scaled limit norm $t_\infty$ is also identified, and since the product $t_\infty \times \beta_\infty$ is identified, it follows that $\beta_\infty$ is identified as well.

### APPENDIX C: SUPPLEMENTARY EXPERIMENTS AND EXPERIMENTAL DETAILS

**C.1. Supplementary experiments.** We describe a sequence of experiments aiming to illustrate the performance of the SVD estimators defined in Section 4. The aim is threefold: first, to show that their empirical behavior is the one predicted by Propositions 1 and 2. Second, to show that they behave similarly to the Bayesian estimators we consider in the main text. Third, we show that the Shrinkage phenomena described in the main text for our Bayesian estimators also replicate in these rate-optimal SVD estimators.

The results of the main experiment are shown in Fig. S1. We study the MSE of the SVD-based estimators for subspaces $\beta', t'$, the estimator of signal strength $\beta_N^2 \times t_D^2$, and the resulting estimator of $\beta$. We consider growing $N, D \in \{5, 10, 20, 50, 100, 250, 500\}$ times are sampled evenly in $[0, 1]$ and that $\beta_0^n = 0, \beta^n \equiv \beta_1^n \sim \mathcal{N}(0, 10)$ (we assume $\beta_0 = 0$ so we don't infer it; and the same applies for the rest of experiments here). We consider an observation noise parameter $\phi^2 = 0.1$. Results show that the MSE of $t'$ and $\beta'$ (average over components) grows to zero at the rate $1/(DN)$ (Fig. S1B-C), as predicted by Proposition 1. However, as shown in S1, the MSE of $\beta$ doesn't converge at this rate, and a big number of features $N$ are necessary to have an MSE comparable to the baseline where times are known. This behavior is consistent with the one described in Fig. 5B in the main text, although the estimators are different. This suggests that the Bayesian estimator may achieve near-optimal performance. The slowdown in convergence of the MSE of $\beta$ is explained by a slower convergence of the signal strength $\bar{\beta}_N^2 \times \bar{t}_D^2$. As suggested by Proposition 2, we may need to make both $N$ and $D$ be extremely large to have an error converging to zero.

Additionally, in Fig. S2 we show examples of inferred pseudotime sequences. We assume that times were sampled from a cubic distribution (same as described in C.3.4), that $\beta^n \sim \mathcal{N}(0, 1)$, $\phi^2 = 0.1$ These results are analog to Fig. 6B (third column) in the main text and show that the SVD-based estimator exhibits similar behavior. Moreover, we show that, as expected, as the number of features $N$ grow, we achieve arbitrary precision in estimation of the sequence of pseudotimes. This idea is at the core of the proof of Proposition 2.

Finally, the experiment in Fig. S3 aims to complement our discussion on the benefits of using Shrinkage estimators. In the main text (Section 5.3) we showed that Shrinkage over $\beta$ had only a modest effect in pseudotime estimation in the linear-Gaussian case, but this effect was much more visible in the negative binomial case (see Fig. 7 and supplementary Figs. S7, S8 and S9). Here, we construct an analog (but different) situation that replicates this phenomenon, this time using the SVD-based analysis. In this case, we assume that $\beta^n = 1$, times are chosen from a cubic sequence, $D = 20$, $N \in \{1, 10, 25, 100, 250, 500\}$. Also, unlike previous examples, we consider the noise distribution $0.5 \times S(\nu)$ where $S(\nu)$ is a Student's t-distribution with $\nu$ degrees of freedom, and $\nu \in \{2, 3, 5, 30, 100\}$, so that if $\nu$ is large we recover the Gaussian case but for small $\nu$ it corresponds to heavy-tailed errors.

We consider the task of estimating the normalized pseudotime sequence $t'$. As estimators, we consider the SVD-based estimator and a structure-aware, 'hierarchical' estimator that takes into account the fact that all $\beta^n$'s are equal (so there is an underlying trivial hierarchical model). We achieve this by taking as estimator of $t'$ simply the normalized quantity

$$\hat{t}_d = \frac{\frac{1}{N}\sum_{n=1}^{N} X_d^n}{||\frac{1}{N}\sum_{n=1}^{N} X_\cdot^n||}.$$

In Fig. S3, we compare both estimators and show that while they have comparable performance for large $\nu$, in the event of heavy-tailed errors (small $\nu$) the hierarchical estimator enjoys a much smaller error. This adds up to the evidence that while pseudotime inference can benefit from shrinkage estimators on $\beta$, such benefits are more likely to manifest beyond the Gaussian-linear case. The fact that we recover this behavior beyond the Bayesian setup suggests that this phenomenon doesn't have to do with our choice of a Bayesian framework.

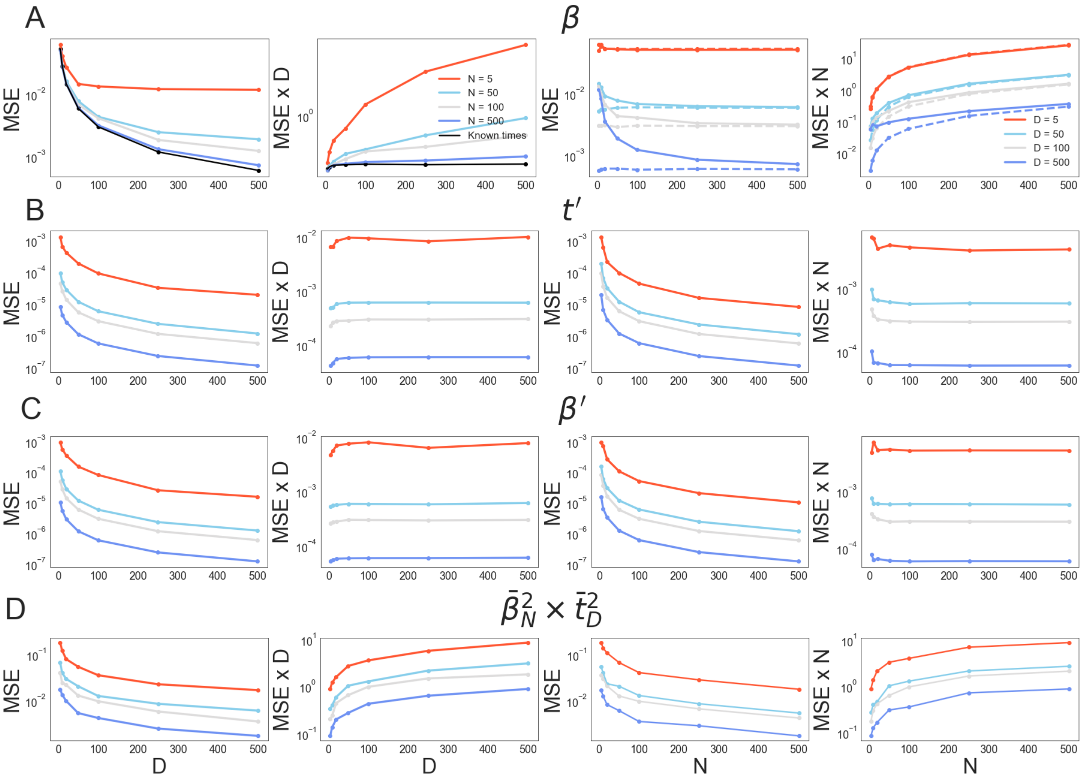

FIG S1. ***Error analysis of for the SVD-based estimators***. *The first column shows the MSE as a function of D. The second column shows the quantity $MSE \times D$ as a function of $D$. Likewise, the third and fourth column show $MSE$ and $MSE \times N$ as a function of $N$. While in all cases the error decreases as $N$ or $D$ increase, from the second and fourth column we can deduce whether the decrease occurs faster than $1/D$ or $1/N$.*

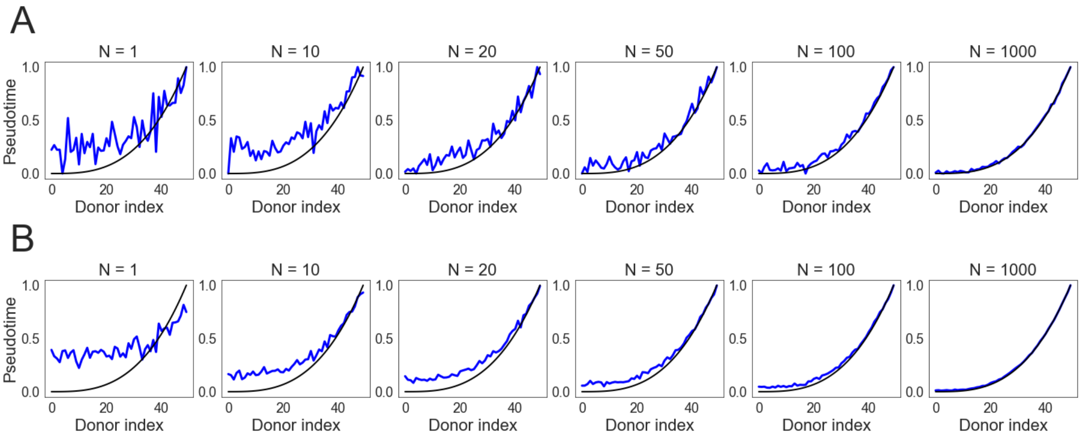

FIG S2. ***Pseudotime inference*** *A: Comparison between inferred pseudo time sequence (in blue, after re-scaling to enforce that minimum and maximum are 0 and 1, respectively) and true pseudotime sequence (black). B: Same as A, but averaging inferred pseudotimes over multiple (1000) experiments.*

**C.2. Supplementary figures.** In Fig. S4 we complement the results of Fig 5A by considering more noise variances $\phi^2$ and by showing the resulting posterior variances of $\beta$ (sum of the variances of $\beta_0$ and $\beta_1$). All results of Fig. S4 correspond to the choice of a uniform prior on pseudotimes and equispaced true times, with a single feature.

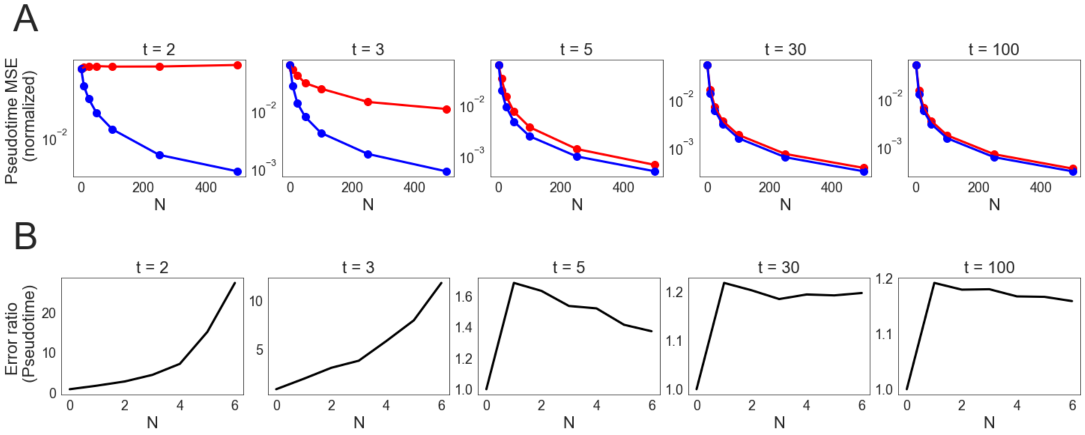

FIG S3. ***Pseudotime error analysis****.A: Comparison of MSE (average over pseudotimes) of the SVD-based estimator (red) and the structure-aware, hierarchical estimator (blue) for the normalized pseudotimes $t'$, as a function of number of features $N$. B: ratio of the error of the SVD-based estimator in relation to the hierarchical estimator. For small $\nu$ the hierarchical estimator vastly outperforms the SVD-based estimator. For large $\nu$ both estimators are indistinguishable in their errors.*

Figures S5 and S6 complement Figs. 5A 5B, respectively, as they include results for the cubic choice of true pseudotime distribution $\mathbf{t}$, not shown in the main text.

Figures S7, S8 and S9 complement Fig. 7 in the main text. While Fig. 7 is based on aggregate experiments across a number of experimental conditions (different donors $D$, feature variances $\sigma^2$ (or $A$ in the negative binomial case) and noise variances $\phi^2$, these supplemental figures give more detailed results once conditioning on particular values of these variables. Fig. S7 focuses on the Gaussian linear model and Fig. S8 focuses on the non-linear negative binomial model, both for varying $\phi^2$ and $\sigma^2$ or $A$. In Fig. S9 we study the role of the number of donors in both models. Altogether, these figures expand upon the message in the main text, i.e. that cross-benefits of shrinkage estimators on pseudotimes were only observed for the negative binomial mode. Additionally, these gains are larger for a low number of donors compared to a number of features.

**C.3. Experimental details.** Here we give details on the experimental setup of Section 5 in the main text and Sections C.1 and C.2 in the Appendix.

C.3.1. *Bayesian setup.* In all cases where Bayesian estimators were involved, we sampled $n_{mcmc} = 500$ samples from the posterior distribution over $t$ and $\beta$ using 2000 warmup samples. For priors over $t$ we used either a $Beta(1,1)$ distribution (i.e. uniform in $[0,1]$) or $Beta(0.2,0.2)$. Regarding $\beta$, the default is to consider a fully factorized model, i.e. an i.i.d. prior of the form

$$\beta_1^n, \beta_0^n \sim \mathcal{N}\left(0, \sigma_\beta^2\right),$$

and $\sigma_\beta^2$ was assigned the extremely large value of 10000 to represent agnosticism. In the hierarchical model (Section 5.3 only) we used the hierarchical priors

$$\beta_i^n, \sim \mathcal{N}\left(\beta_i, \sigma^2\right), \beta_i \sim \mathcal{N}\left(0, \sigma_\beta^2\right).$$

Again, $\sigma_\beta^2$ was chosen very large. Regarding $\sigma^2$, this value was either assumed to be known or sampled from its own prior distribution ($HN$ denotes the half-normal distribution):

$$\sigma^2 \sim HN(500).$$

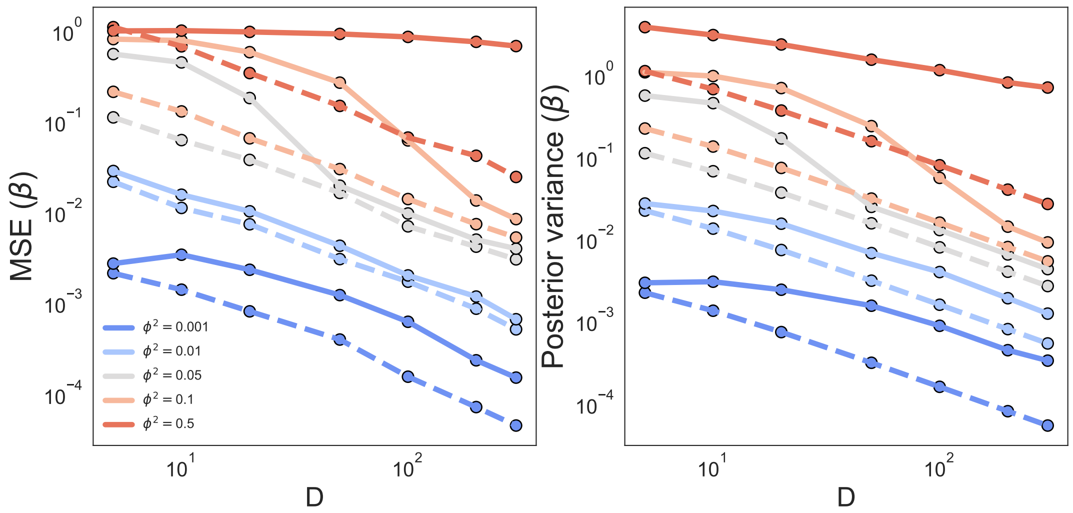

FIG S4. ***Elementary convergence study for the linear model (Eq.** (4)**) with a single** ($N = 1$) **feature**. Left: MSE of $\beta = (\beta_0, \beta_1)$ for different noise variances $\phi^2$ (colored lines) and number of sampled pseudotimes ($D$, x-axis). Dashed lines correspond to the baseline where $t_d$'s are known (i.e., solving a linear regression). Right: sum of posterior variances of $\beta_0$ and $\beta_1$.*

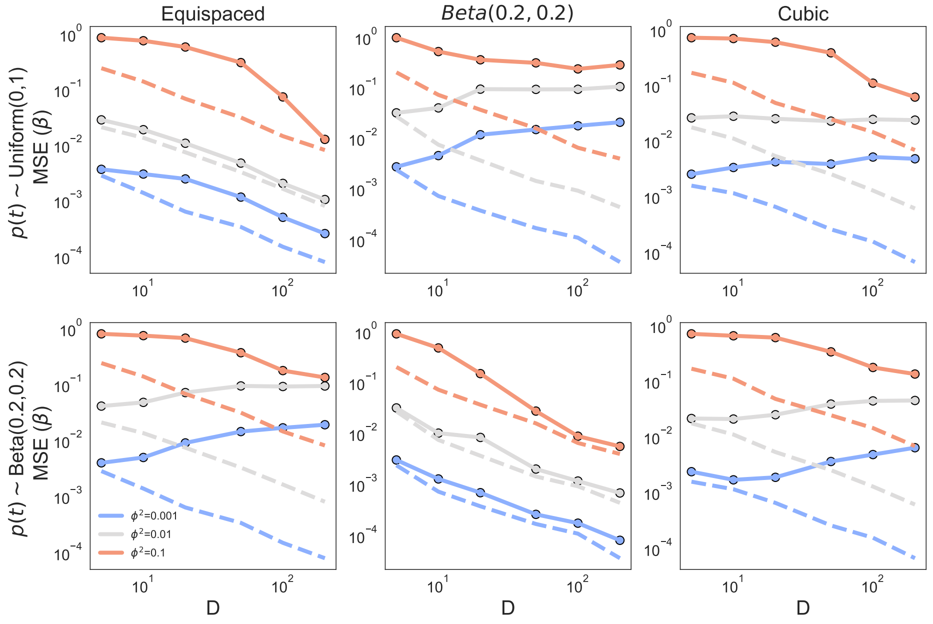

FIG S5. ***Effect of choices of pseudotime priors in the linear model (Eq.** (4)**) on estimation error (MSE) of** $\beta = (\beta_0, \beta_1)$. Priors $p(t) \sim Uniform(0, 1)$ and $p(t) \sim Beta(0.1, 0.1)$ are shown on different rows and population distribution of pseudotimes **t** on different columns. Colors indicate noise levels $\phi^2$. We used a single feature for this experiment ($N = 1$). Dashed lines correspond to the baseline where pseudotimes are known.*

C.3.2. *Error measures.* Given the output of each MCMC run characterized by samples from the posteriors over $t$ and $\beta$, we defined the estimators $\hat{t}_d$ and $\hat{\beta}_i^n$ as the **posterior means**

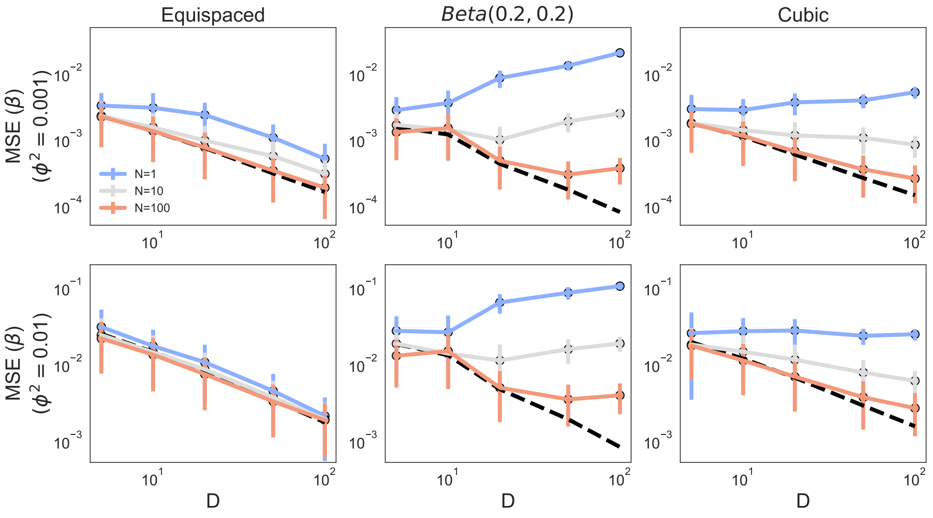

FIG S6. ***Effect of choices of number of features*** $N$ ***and in the linear model (Eq.*** (4)***) on estimation error (MSE) of*** $\beta = (\beta_0, \beta_1)$. *Different rows correspond to different observation noise levels* $\phi^2$*, and columns indicate the true distribution of times* ***t***. *Colors indicate the number of features* $N$. *Features are equal. Specifically, we used* $\beta_0^n = 0$ *and* $\beta_0^n = 1$ *for each* $n \in [1, N]$. *Also, dashed lines correspond to the baseline where pseudotimes are known.*

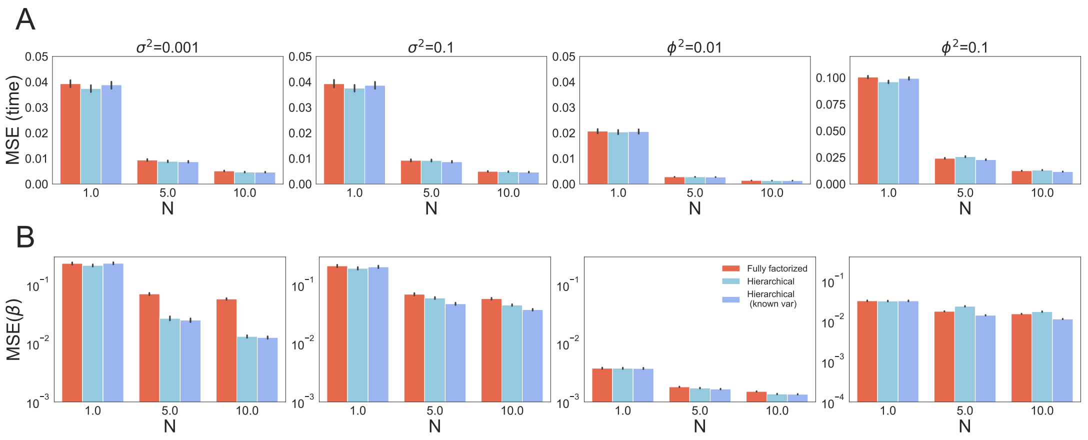

FIG S7. ***Estimation error of different hierarchical models for Eq.*** (4) ***in the linear-Gaussian model***. *The first and second rows correspond to pseudotime and* $\beta$ *errors, respectively. We plot different values of* $N$ *(x-axis), observation noise variances* $\phi^2$*, and feature noise variance* $\sigma^2$.

over these $n_{mcmc}$ samples (but see C.3.3 for details). We analyze these estimators from a frequentist perspective through their MSE. Specifically, we define

$$MSE(\hat{\beta}_i^n) \equiv E\left(\hat{\beta}_i^n - \beta_i^n\right),$$

and

$$MSE(\hat{t}_d) \equiv E\left(\hat{t}^d - t_d\right).$$

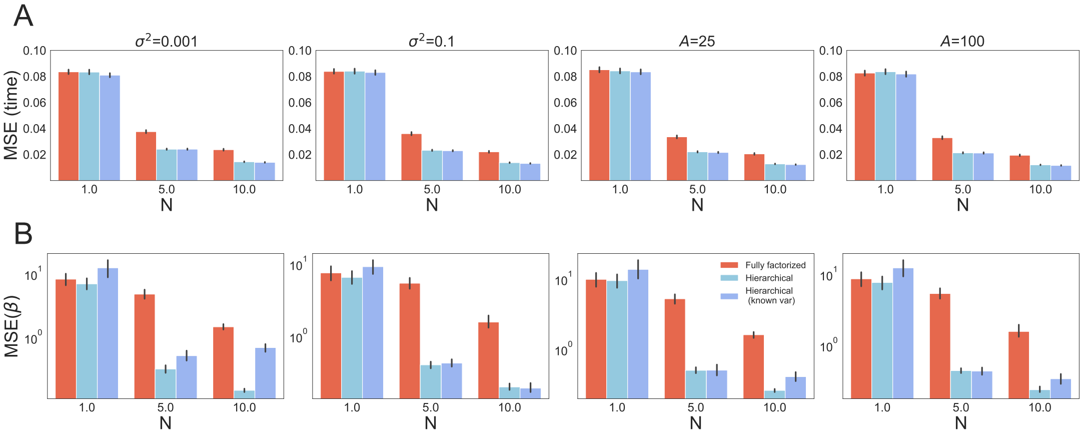

FIG S8. ***Estimation error of different hierarchical models for Eq.*** (4) ***in the negative binomial with logarithmic link function model****. The first and second rows correspond to pseudotime and $\beta$ errors, respectively. We plot different values of $N$ (x-axis), observation noise variances $A$, and feature noise variance $\sigma^2$.*

When we report the overall MSE over $\beta_i$ and $t$ we mean, unless otherwise specified, their average quantities over the entire vectors, i.e.

$$MSE(\hat{\beta}_i) \equiv \frac{1}{N}\sum_{n} MSE(\hat{\beta}_i^n), \quad MSE(\hat{t}) \equiv \frac{1}{D}\sum_{d=1} MSE(\hat{t}_d).$$

Additionally, when we report the $MSE$ of $\hat{\beta}$ we mean the sum over $\hat{\beta}_1$ and $\hat{\beta}_2$,

$$MSE(\hat{\beta}) \equiv MSE(\hat{\beta}_0) + MSE(\hat{\beta}_1).$$

Finally, as all the above MSE are simulated population quantities, we simulate them These quantities are estimated from experiments. Specifically, we run a number $n_{exp} = 200$ of experiments with the same experimental conditions. We estimated the above $MSE$ as the average squared error over those simulations. Whenever we display errorbars in our plot those corresponds to $95\%$ confidence intervals derived from those simulations.

C.3.3. *Re-scaling.* As mentioned above, our estimators $\hat{\beta}$ and $\hat{t}$ are posterior means over simulations. However, we normalize estimators $\hat{t}$ to impose a support constraint. Before averaging we stretch each vector $t$ sampled from the posterior distribution to enforce that its minimum value is 0 and its maximum is 1. Then, we normalize the corresponding $\beta$ samples so that the quantity $\beta_0 + \beta_1 t$ is preserved.

This normalization is particularly relevant in low signal to noise regimes or low $N$ regimes. In those case identifiability is not guaranteed, and in practice the estimated times may appear uniformly close to $0.5$, exhibiting low variability. Then, this heuristic aims to enhance identifiability of our Bayesian estimators. Additionally, we considered the above normalization procedure but after computing the posterior mean of the estimators. We did not observe substantial differences between both.

C.3.4. *Experimental parameters.* In the experiments corresponding to Fig. 5, and Figs. S4, S5 and S6 we used the following parameters

- $D \in \{5, 10, 20, 50, 100, 200, 300\}$
- $\phi^2 \in \{0.001, 0.01, 0.05, 0.1, 0.5\}$
- $N \in \{1, 10, 100\}$

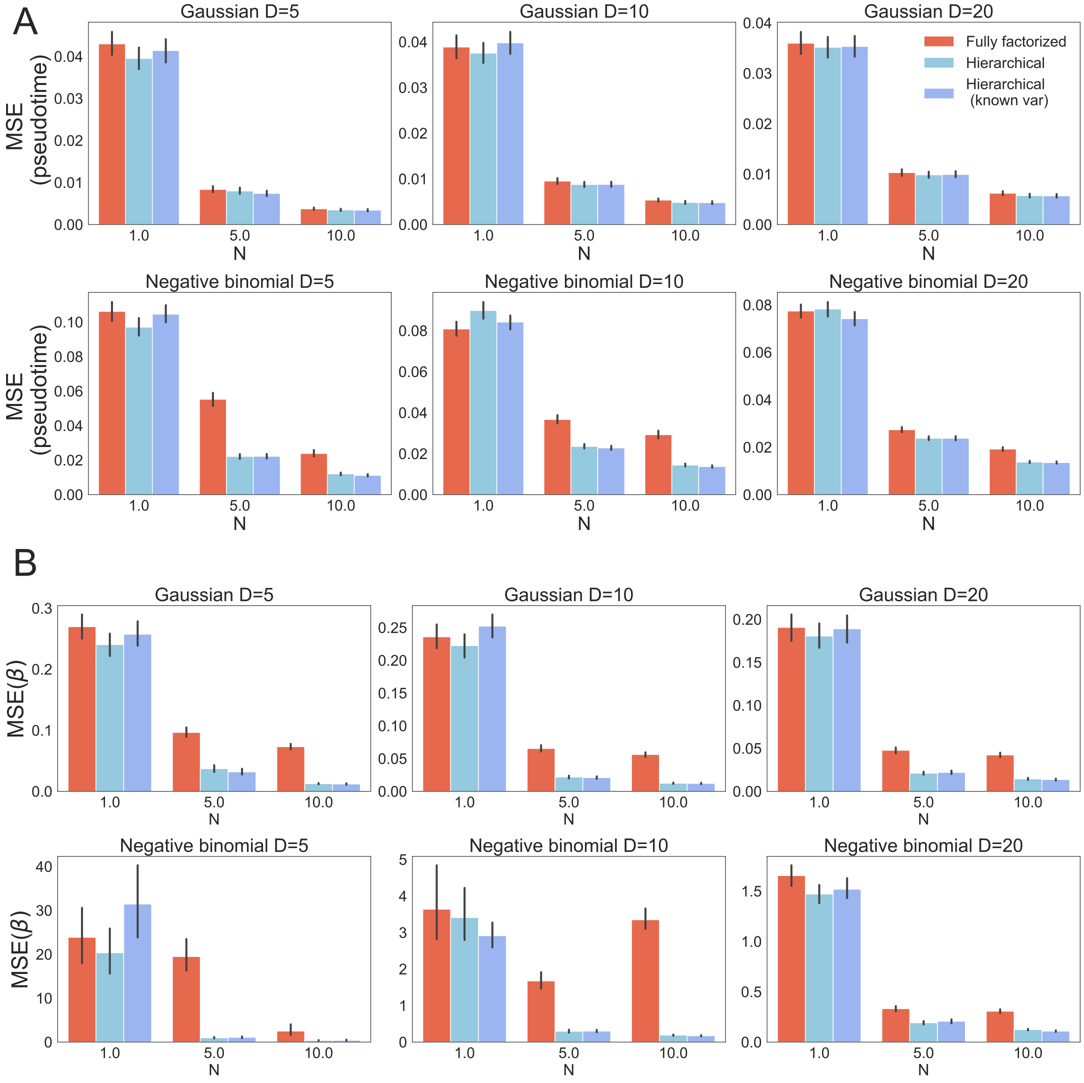

FIG S9. ***Estimation error of pseudotime (A) and $\beta$ (B) of different hierarchical models for Eq.*** (4) ***in the Gaussian (first rows) and negative binomial with logarithmic link function model (second rows)***. *We plot different values of $N$ (x-axis), and varying number of donors $D$*

- $\mathbf{t}$ chosen from i) equispaced grid of $[0, 1]$, ii) sampled from $Beta(0.2, 0.2)$ and iii) chosen from the normalized sequence $t^3$ (enforcing that $\min t_d = 0, \max t_d = 1$).
- $\beta_0^n = 0, \beta_1^n = 1$

In the experiments corresponding to Fig. 6 we used the parameters

- $D \in \{5, 10, 20, 50, 100\}$
- $\phi^2 \in \{0.001, 0.01, 0.1\}$
- $N = 10$
- $\sigma^2 \in \{0, 10\}$
- $\mathbf{t}$ chosen from i) equispaced grid of $[0, 1]$, ii) sampled from $Beta(0.2, 0.2)$ and iii) chosen from the normalized sequence $t^3$ (enforcing that $\min t_d = 0, \max t_d = 1$).
- $\beta_0^n = 0, \beta_1^n \sim \mathcal{N}(0, \sigma^2)$.
- Uniform prior on times.

In the experiments corresponding to Fig. 7 we used

- $D \in \{5, 10, 20\}$
- $\phi^2 \in \{0.001, 0.01, 0.1\}$ (Gaussian case) and $A \in \{10, 25, 100\}$ (negative binomial case)
- $N \in \{1, 5, 10\}$
- $\sigma^2 \in \{0, 0.1, 0.5\}$
- $\mathbf{t}$ chosen from i) equispaced grid of $[0, 1]$, ii) sampled from $Beta(0.2, 0.2)$ and iii) chosen from the normalized sequence $t^3$ (enforcing that $\min t_d = 0, \max t_d = 1$).
- Uniform prior on times.
- $\beta_0^n \sim \mathcal{N}(0, \sigma^2), \beta_1^n \sim \mathcal{N}(1, \sigma^2)$ (Gaussian case) and $\beta_0^n \sim \mathcal{N}(0, \sigma^2), \beta_1^n \sim \mathcal{N}(2, \sigma^2)$ (negative binomial case).

For the results in Fig. 8 we used the parameters.

- $D = 50$,
- $A = 100$ (negative binomial case)
- $N = 2$
- $\mathbf{t}$ chosen from an equispaced grid.
- $\beta_0^n = 0, \beta_1^n \sim \mathcal{N}(0, \sigma^2)$.
- Uniform prior on times.
- $\beta_0^1 = 4.0, \beta_1^1 = -4.0, \beta_0^2 = 0$, $\beta_1^2 \in \{2, 4, 6\}$.

Sampled datasets $X_d^n$ corresponding to different choices of $\beta_1^2$ are shown in Fig. S10. The symmetric case corresponds to $\beta_1^2 = 4$, where each channel contributes a comparable amount of information.

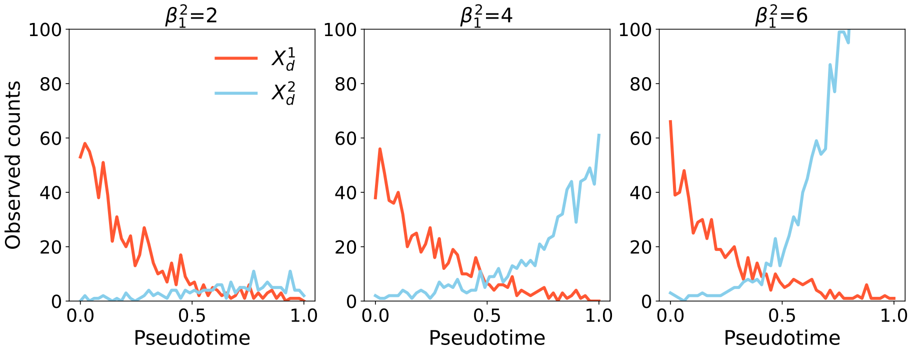

FIG S10. ***Example of sampled count sequences in the setup of Fig. 8*** *. Counts in the first channel (red) are always sampled with parameters* $\beta_0^1 = 4, beta_1^1 = -4$. *In the second channel (light blue) the parameters are* $\beta_0^2 = 0$ *and* $\beta_1^2 \in \{2, 4, 6\}$ *(different plots).*

**C.4. Supplementary figures for fits with neuropathological data.** The following section consists of figures that have been referred to in the main text in the section 6.

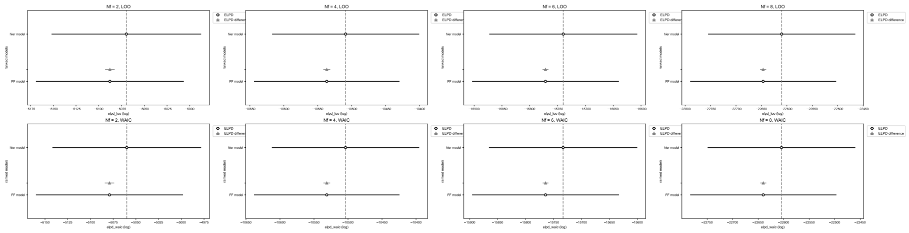

FIG S11. ***Model comparison using ELPD**. Comparing ELPD of the two bayesian inference models with leave-one-out cross validation (LOO, top row) and WAIC criterion (bottom row) across* $M = 2 - 8$ *features. While both models perform comparably, the hierarchical model performs slightly better*

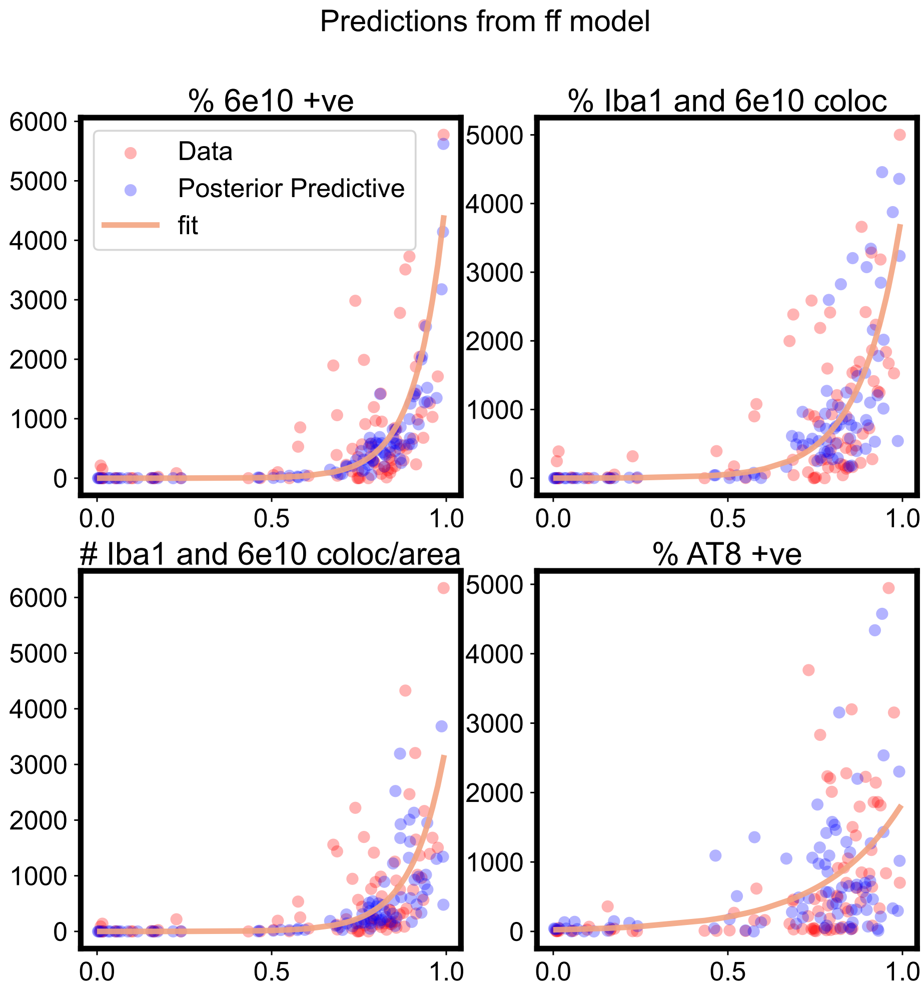

FIG S12. ***Visualization of modeling predictions.*** *Model results overlaying the pathology data for listed feature (red dots) with model's posterior predictive (blue dots). The solid curve is the mean prediction from the deterministic model. The results are plotted for 4 example pathological features.*

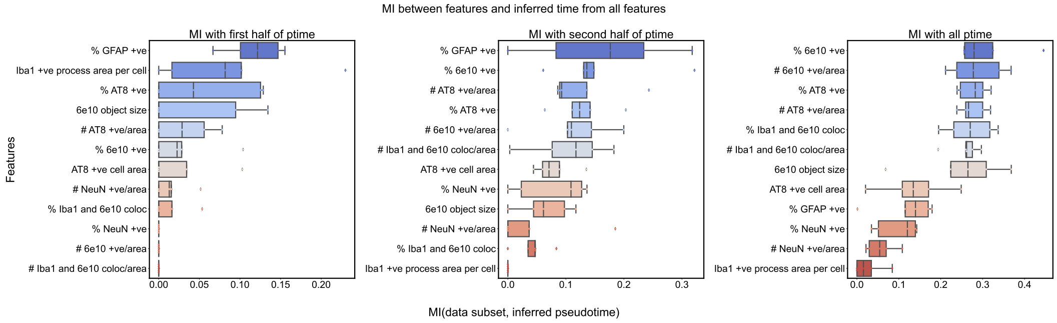

FIG S13. ***Mutual Information shows reliance on different features for different parts of the pseudotime inference.*** *Left: Mutual Information (MI) is calculated between early pseudotimes (0-0.5) inferred from all important pathological features and individual pathological feature. Middle: MI calculated between late pseudotime (0.5-1) and pathological features. Right: MI calculated between all pseudotime (0-1) and features. Since most donors lie in the late pseudotime regime, the features that are informative for late pseudotime are also dominant in the overall pseudotime inference.*

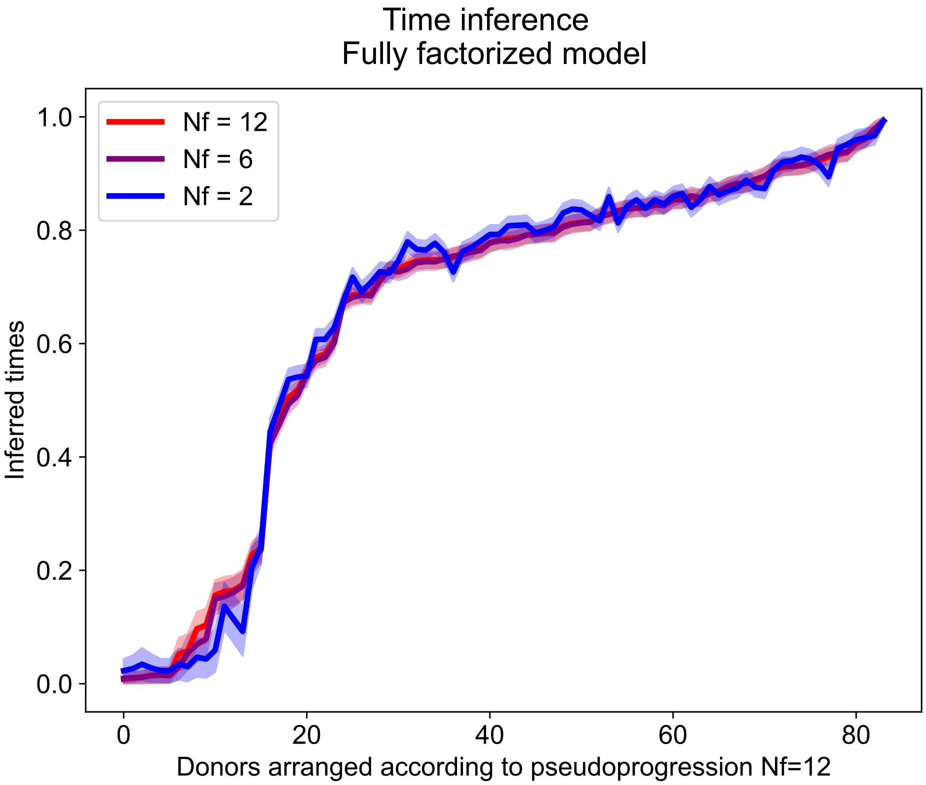

FIG S14. ***Comparison of pseudotime shows decreasing impact of increasing number of features in the bayesian inference model****. From blue to red, the number of features utilized for pseudotime inference is increased. However, beyond 6 features, there is not much impact on increasing feature data on pseudotime inference, either in terms of donor ordering through mean pseudotime or decreasing the error in pseudotime inference (shaded region).*

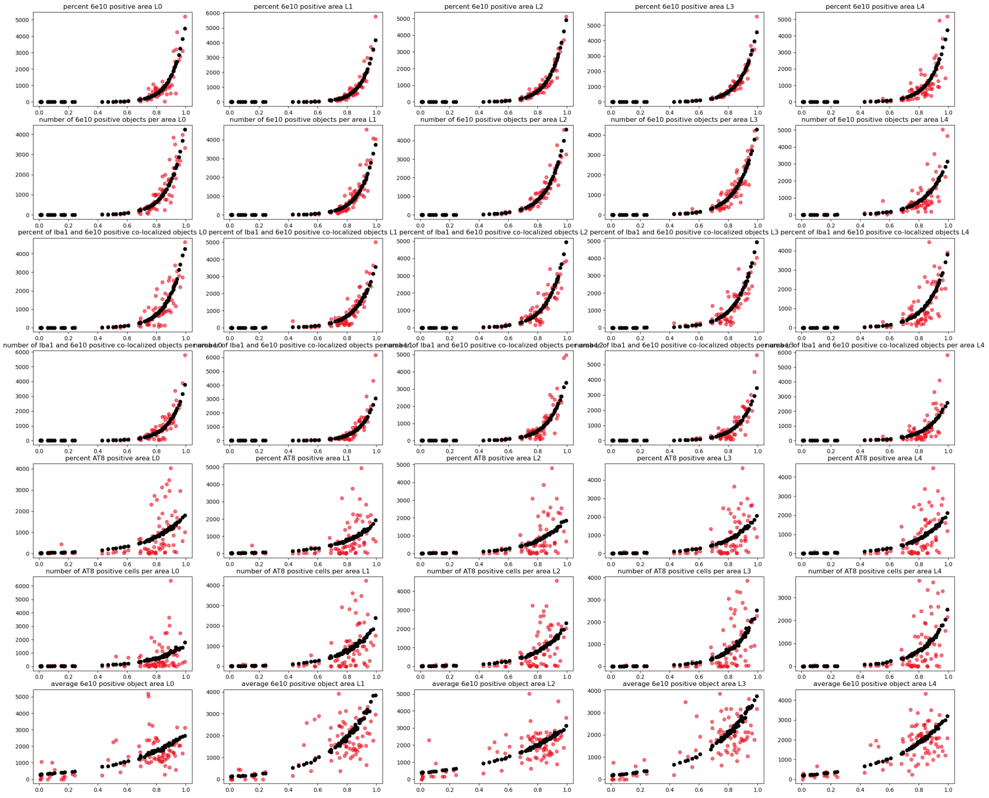

FIG S15. ***Fits from minimal model including staging information show reasonable agreement with data.*** *Left to right: Data from different cortical layers of the same pathological features. Top to bottom: Data from the 7 pathological markers used in the minimal model with staging information.*

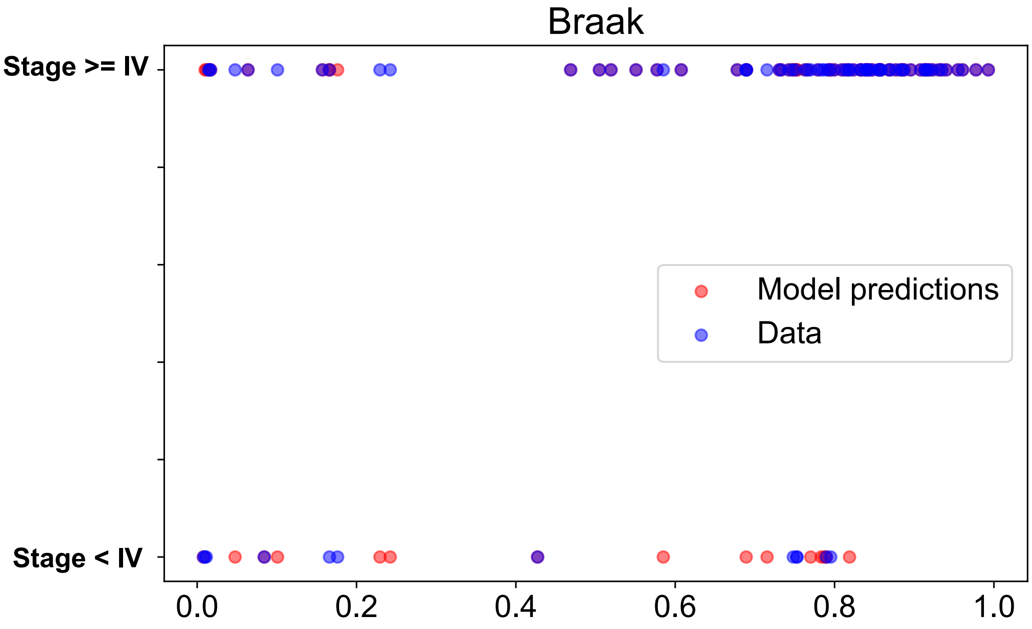

FIG S16. *Fitting a binarized version of braak staging information (Stage >=IV = 1, Stage < IV = 0) from minimal model including staging information. The Spearman correlation of the binarized Braak stages with our predicted pseudotime is* 0.35. *F1 score for the model with staging predictions = 0.86*
